# Structures of PSI-FCPI from *Thalassiosira pseudonana* in high light provide convergent evolution and light-adaptive strategies in diatom FCPIs

**DOI:** 10.1101/2024.05.30.596378

**Authors:** Yue Feng, Zhenhua Li, Yang Yang, Lili Shen, Xiaoyi Li, Xueyang Liu, Xiaofei Zhang, Jinyang Zhang, Fei Ren, Yuan Wang, Cheng Liu, Guangye Han, Xuchu Wang, Tingyun Kuang, Jian-Ren Shen, Wenda Wang

## Abstract

Diatoms achieve great survival success in the fluctuating oceanic environment, rely on fucoxanthin chlorophyll *a*/*c*-binding proteins (FCPs) to complete light harvesting and quenching, which provide about 20% primary productivity on earth. We report two cryo-electron microscopic structures of photosystem I (PSI) with 13 or 5 FCPIs respectively at 2.78 Å and 3.20 Å resolution from *Thalassiosira pseudonana* under high light conditions. 8 Lhcr FCPIs are found detached from the PSI-13FCPI supercomplex under high light conditions, remaining 5 FCPIs are stably combined with the PSI core including Lhcr3, RedCAP, Lhcq8, Lhcf10, and FCP3 subunits. The specific pigment network in this centric diatom *T. pseudonana* demonstrates a higher proportion of Chlorophylls *a*, diadinoxanthins, and diatoxanthins but fewer fucoxanthins compared with the huge PSI-FCPI from another centric diatom *Chaetoceros gracilis*, thus exhibiting more efficiency in energy transfer and dissipation among FCPI antennas. These results reveal the assembly mechanism of several types of peripheral FCPIs and corresponding light-adaptive strategies in *T. pseudonana*, as well as the convergent evolution of the diatoms PSI-FCPI structures.

## Introduction

Photosynthesis converts solar energy into chemical energy and releases oxygen, light energy is absorbed by the peripheral antennas and transferred to the two photosystems (photosystem I and photosystem II, PSI and PSII)^1^. In PSI, the efficiency of charge separation and photosynthetic electron energy transfer is close to 100%^2,3^.

Eukaryotic photosynthetic organisms can be classified into green and red lineages^4^, the PSI cores of green lineages are conserved among green algae^5–8^, mosses^9–11^ and higher plants^2,12,13^, and the number of bound LHCIs is correlated with their evolutionary status and environment^6^. In the case of light quality changes and treatment by NaN3/NaF, some phosphorylated LHCIIs move laterally from PSII to PSI leading to state transition and forming the unique PSI-LHCI-LHCII supercomplex^14–16^.

In red lineages such as red algae^17–19^, cryptophytes^20^ and diatoms^21,22^, most subunits in PSI core are also conserved, but different species have differences in the combination of some subunits such as PsaR, PsaK, PsaS and PsaO^23^, thus affecting the number of peripheral antennas.

Diatoms are important red lineage phytoplanktons and provide around 20% of primary productivity on earth^24^. Diatoms rely on unique fucoxanthin chlorophyll *a*/*c* binding proteins (FCPs) to harvest light energy, fucoxanthin (Fx) and chlorophyll *c* (Chl *c*) expert in blue-green light harvesting and efficiently excited energy transfer^25^. In addition, diatoms dissipate excess light energy with the help of the diadinoxanthin (Ddx)/diatoxanthin (Dtx) cycle^26^, The protein families of diatom antennas are much more than plants^27,28^, Lhcf and Lhcr mainly as antennas in the periphery of PSII and PSI, respectively^21,22,29^. Lhcx involved in the light-stress response^30^, Lhcq subfamily mainly make up the second FCPIs belt^22,31^, some special Lhcr were categorized to a small Lhcz subfamily^31^. Furthermore, a suspected RedCAP (red lineage chlorophyll *a*/*b*-binding-like protein) family protein (fc13194 in the diatom *Fragilariopsis cylindrus*) was preliminary assigned to FCPI-1 due to the broken local density map in PSI-FCPI of *Chaetoceros gracilis*^21^.

Previous studies have demonstrated that the PSI core of diatoms can bind 16 to 24 FCPIs in *C.gracilis* under low-light conditions^21,22^. Recently, the structural analysis of PSII-FCPII from *Ditylum* (*C.gracilis*) and *Coscinodiscaceae* (*Cyclotella meneghiniana* and *Thalassiosira pseudonana*) has demonstrated that centric diatoms have structural heterogeneity in PSII supercomplex^32–34^. However, the structural models at 17 Å resolution by EM and Cryo-ET also proved that PSI-FCPI supercomplex from diatom *T.pseudonana* is structurally convergent with the Cg-PSI-FCPI complexes^35^. On the other hand, relative abundances of FCPI composition prominent change on high light effect in Tp-PSI-FCPI^36^. The phylogenetic relationship between *C.gracilis* and *T.pseudonana* has been elucidated^31^. Thus, resolving the Tp-PSI-FCPI under high light will provide more information about different FCPIs adaptive to light effects and further validate the convergent evolution of PSI in diatom.

In this study, we resolved two structures of PSI-FCPI supercomplex under high light from *T.pseudonana* by Cryo-EM, with either 13 or 5 FCPIs associated with the PSI core, at an overall resolution of 2.78 Å and 3.20 Å, respectively. The results provide detailed information on the protein and pigments arrangement of Tp-PSI-FCPI under high light, most FCPIs in PSI-13FCPI conserved compared with Cg-PSI-FCPI demonstrating that different PSI-FCPI show convergent evolution in diatoms. Structures between PSI-13FCPI and PSI-5FCPI provide insights into the light-adaptive strategy and potential assembly mechanism in diatom PSI-FCPI.

## Results

### Characterization of Tp-PSI-FCPI in low light and high light

The PSI-FCPI supercomplexes were purified from *T.pseudonana* (Tp) cells grown under low-light (LL) and high-light (HL) conditions by sucrose density gradient ultracentrifugation and characterized by SDS–polyacrylamide gel electrophoresis (Fig. 1a, b) (see Methods for more details). High-performance liquid chromatography(HPLC) analysis indicates that both Tp-PSI-FCPI-LL and Tp-PSI-FCPI-HL contain six pigments including Chl *a*, Chl *c*, Fx, Ddx, Dtx and β-carotene (Bcr). In addition, the ratio of Dtx/Ddx is higher in the Tp-PSI-FCPI-HL than Tp-PSI-FCPI-LL, and other pigments almost same between two light conditions (Fig. 1c). We further purified and characterized Tp-PSI-FCPI-LL and Tp-PSI-FCPI-HL by gel filtration chromatography, among which one major peak elution and two small peak elutions were found in PSI-FCPI-LL, we named them PSI-FCPI-Huge (11.7 mL), PSI-FCPI-Large (PSI-FCPI-L, 13.1 mL), PSI-FCPI-Small (PSI-FCPI-S, 14.5 mL) according to their molecular weight, two similar peak elutions were found in PSI-FCPI-HL, but the content of PSI-FCPI-S increased apparently (Fig. 1d).

**Fig. 1.**
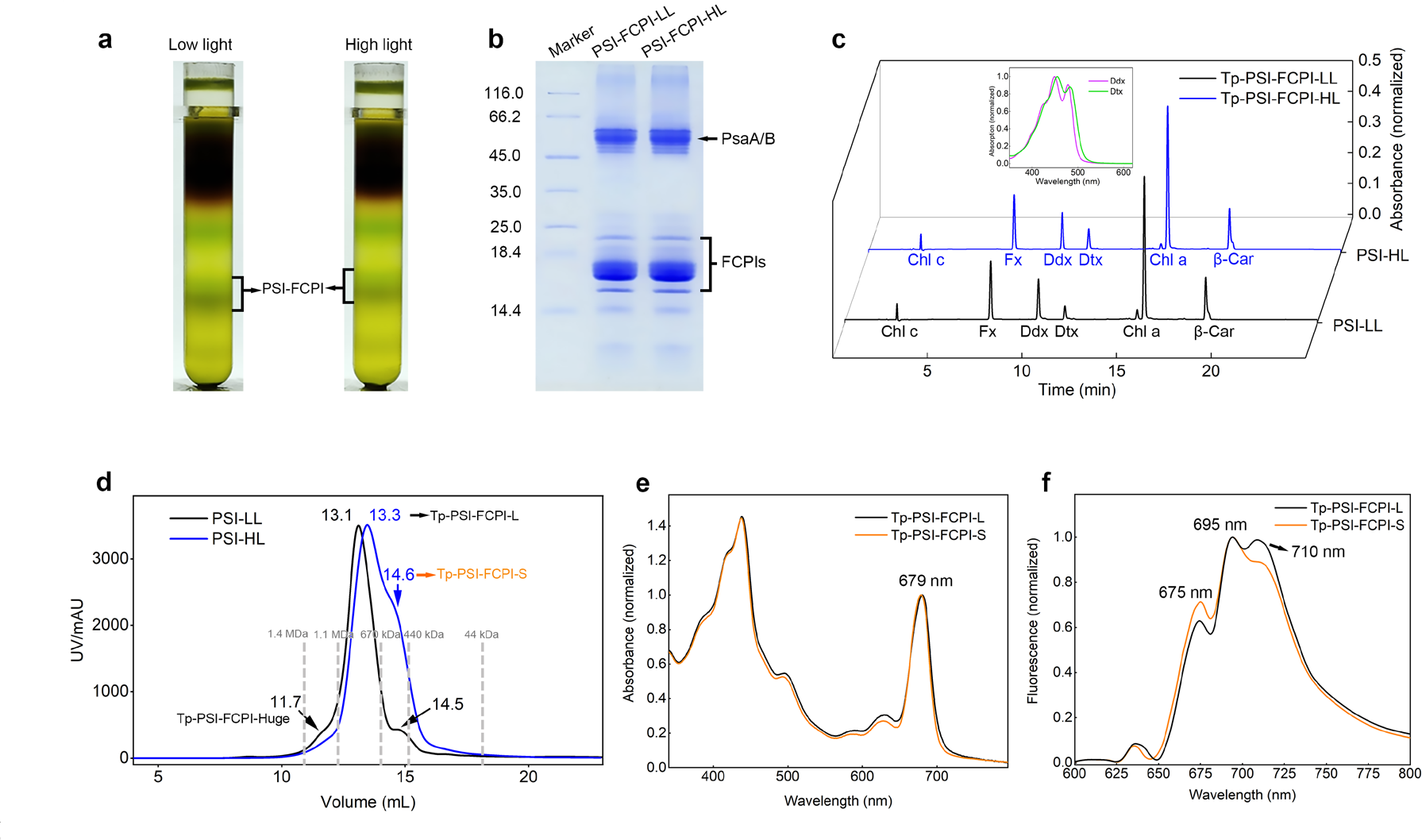
**Purification and characterization of PSI-FCPI from *T. pseudonana*. a**, Isolation of PSI-FCPI by sucrose density gradient centrifugation. The FCPI-FCPI bands were labeled under different light conditions. **b**, SDS-PAGE analysis of the purified PSI-FCPI from *T. pseudonana*. Lane 1: molecular weight marker (Thermo Fisher Scientific: 26610); Lane 2: purified PSI-FCPI under low light (5 μg Chl); Lane 3: purified PSI-FCPI under high light (5 μg Chl). **c**, Analysis of the pigments of PSI-FCPI purified from diatom cells under low light and high light by HPLC monitored at 445 nm and normalized based on the content of Chl *a*. Six major pigment peaks were identified, including Chl *a*, Chl *c*, Fx, Ddx, Dtx, and Bcr. **d**, Normalized elution profile of the PSI-FCPI supercomplex under low light and high light by size-exclusion chromatography (Cytiva, Superose 6 Increase 10/300 GL) at the absorbance of 280 nm, respectively. The gray dashed line is the peak of size exclusion chromatography of the high molecular weight standard samples. **e**, Room temperature absorption spectrum of PSI-FCPI-L and PSI-FCPI-S. **f**, Normalized 77 K fluorescence emission spectra of PSI- FCPI-L and PSI-FCPI-S excited at 436 nm.

The Cryo-EM structures of huge PSI-FCPI supercomplex from *C.gracilis* at low light have been solved into Cg-PSI-16FCPI and Cg-PSI-24FCPI, respectively^21,22^. The PSI supercomplex in present centric diatom *T.pseudonana* was suspected to combine with 18 FCPIs^35^, which may correspond to our small 11.7 mL peak elution in Tp-PSI-FCPI-LL (Fig. 1d), but this component is little.

In this study, we collected the mixed elutions (13 ml-15 mL) of Tp-PSI-FCPI-HL for Cryo-EM and analyse the related light-adaptive strategies of different PSI-FCPI particles. The peak elution of Large at PSI-FCPI 13.3 mL (PSI-FCPI-L) and the shoulder elution of Small PSI-FCPI-S at 14.6 mL (PSI-FCPI-S) were collected for spectroscopic analysis (Fig. 1e, f). The Room temperature absorption spectroscopic results indicate that PSI-FCPI-L has more wavelength region of Fx absorption than PSI-FCPI-S and the 77 K fluorescence emission spectroscopic results show the PSI-FCPI- L has a higher peak of the PSI core in 710 nm (Fig. 1e, f).

### The overall structure of two Tp-PSI-FCPI

The PSI-FCPI samples under high light were imaged by single-particle cryo-EM at 300 kV. 129,745 particles and 53,013 particles were finally used to construct the structural model of Tp-PSI-FCPI-L and Tp-PSI-FCPI-S under high light at 2.78 Å and 3.20 Å resolution, respectively (Supplementary Fig. 1). We construct the structural models of PSI-FCPI-L and PSI-FCPI-S under high light by COOT^37^ and Phenix^38^ (Supplementary Table 1). Based on the HPLC results and density maps obtained in this study and the complete genome data^27^, amino acids, pigments and cofactors were assigned to Tp-PSI-FCPI-L and Tp-PSI-FCPI-S models (Supplementary Fig. 2 and Table 2). Both Tp-PSI-FCPI-L and Tp-PSI-FCPI-S are monomers, PSI core combines 13 FCPIs including Lhcr1、Lhcr3、Lhcr4、Lhcr7、Lhcr10、Lhcr11、Lhcr14、 Lhcr18、Lhcr20、Lhcf10、Lhcq8、FCP3 and RedCAP with an overall molecular weight of 725 kDa (Fig. 2a), 8 Lhcr family proteins were detached in Tp-PSI-FCPI-S, only retain 5 different family FCPIs, Lhcr3、RedCAP、Lhcq8 and Lhcf10, resulting in an overall molecular weight of 510 kDa (Fig. 2b). In addition to the protein subunits, 229/142 Chls *a*, 15/4 Chls *c*, 38/17 Fxs, 27/8 Ddxs, 1 Dtx, 21 Bcrs, 2 plastoquinones, 3 Fe4S4 clusters, 30/19 lipids are assigned to Tp-PSI-FCPI-L or Tp-PSI-FCPI-S, respectively (Supplementary Table 3).

**Fig. 2.**
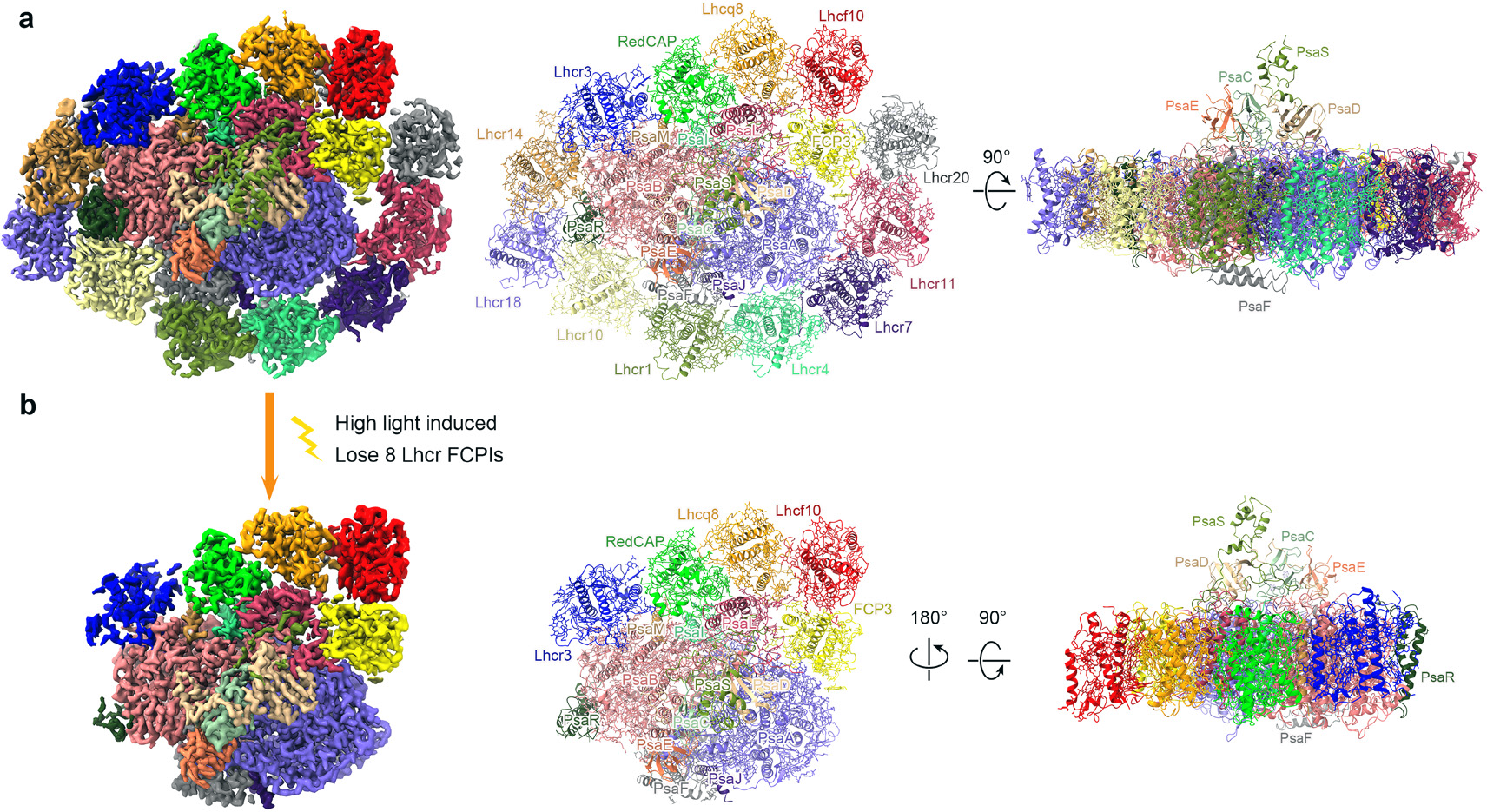
**Overall structure of the PSI-PCPI-L and PSI-FCPI-S from *T. pseudonana*. a**, Cryo-EM density map and overall structure of the PSI-FCPI-L with a top view and a side view, respectively. Each subunit is represented as ribbons and colored differently. **b**, Cryo-EM density map and overall structure of the PSI-FCPI-S with a top view and a side view, respectively. Each subunit is represented as ribbons and colored the same as in (a). The colors of the FCPI subunits are the same as in (a).

The PSI core contains 2 large transmembrane subunits (PsaA, PsaB), 6 small transmembrane subunits (PsaF, PsaI, PsaJ, PsaL, PsaM, PsaR) and 4 extrinsic subunits (PsaC, PsaD, PsaE, PsaS) attached at the stromal surface for efficient electron transfer (Fig. 2). These subunits are conserved with the Cg-PSI core but longer PsaL subunit (Supplementary Fig. 3a). Compared with typical red lineage organisms such as red algae *Porphyridium purpureum*^18^ or cryptophyte algae *Chroomonas placoidea*^20^, the Tp-PSI core loss the PsaK and PsaO. Compared with PSI-LHCI in green lineage organisms, the Tp-PSI core loss PsaG/H/K/O, whose locations were occupied by Tp- PsaR, Tp-Lhcq8, Tp-Lhcr11 and Tp-FCP3 in Tp-PSI-FCPI, respectively (Supplementary Fig. 3b). In addition, the unique PsaS in diatom is modeled in this study based on the sequence of the genome from Val47 to T177 (UniProt B8BUW3)^27^ (Supplementary Fig. 2a and Table 3), which was found in Cg-PSI-FCPI but modeled by poly-alanines due to the limited transcriptome data^21^.

Notably, the FCPI site in Tp-Lhcf10 has a big shift at 6.8 Å compared with Cg- Lhcf3 in conserved diatom PSI-FCPI supercomplexes (Supplementary Fig. 3a), the positions of Tp-Lhcf10 and Cg-Lhcf3 are overlapped with mobile LHCII in the green alga PSI-LHCI-LHCII (Supplementary Fig. 3b), but we did not found any phosphorylation site surround Tp-Lhcf10 in the PSI-FCPI-HL by checking the density map in 2.78 Å, which also indicates that diatoms may not have state transitions^39^. In addition, a predicted RedCAP ortholog Cg-FCPI-1 was not well resolved for their limited local density map^21,23^, the FCPI site in Tp-RedCAP was well-solved in this study for its area has a better density map, we built the model of the N-terminus, unique AC-Helix and assigned all ligands for Tp-RedCAP (Fig. 3 and Supplementary Fig. 2b- c), the FCPI site of Tp-Lhcq8 between Tp-Lhcf10 and Tp-RedCAP was also well modeled (Fig. 3 and Supplementary Fig. 2b-c). The three FCPIs above combined stably in Tp-PSI-FCPI-L/S also exist in Cg-PSI-24FCPI^21^ but lost in Cg-PSI-16FCPI^22^, which may be caused by different growth temperatures, concentrations of CO2^40^ and different detergent treatments to thylakoid^41,42^.

**Fig. 3.**
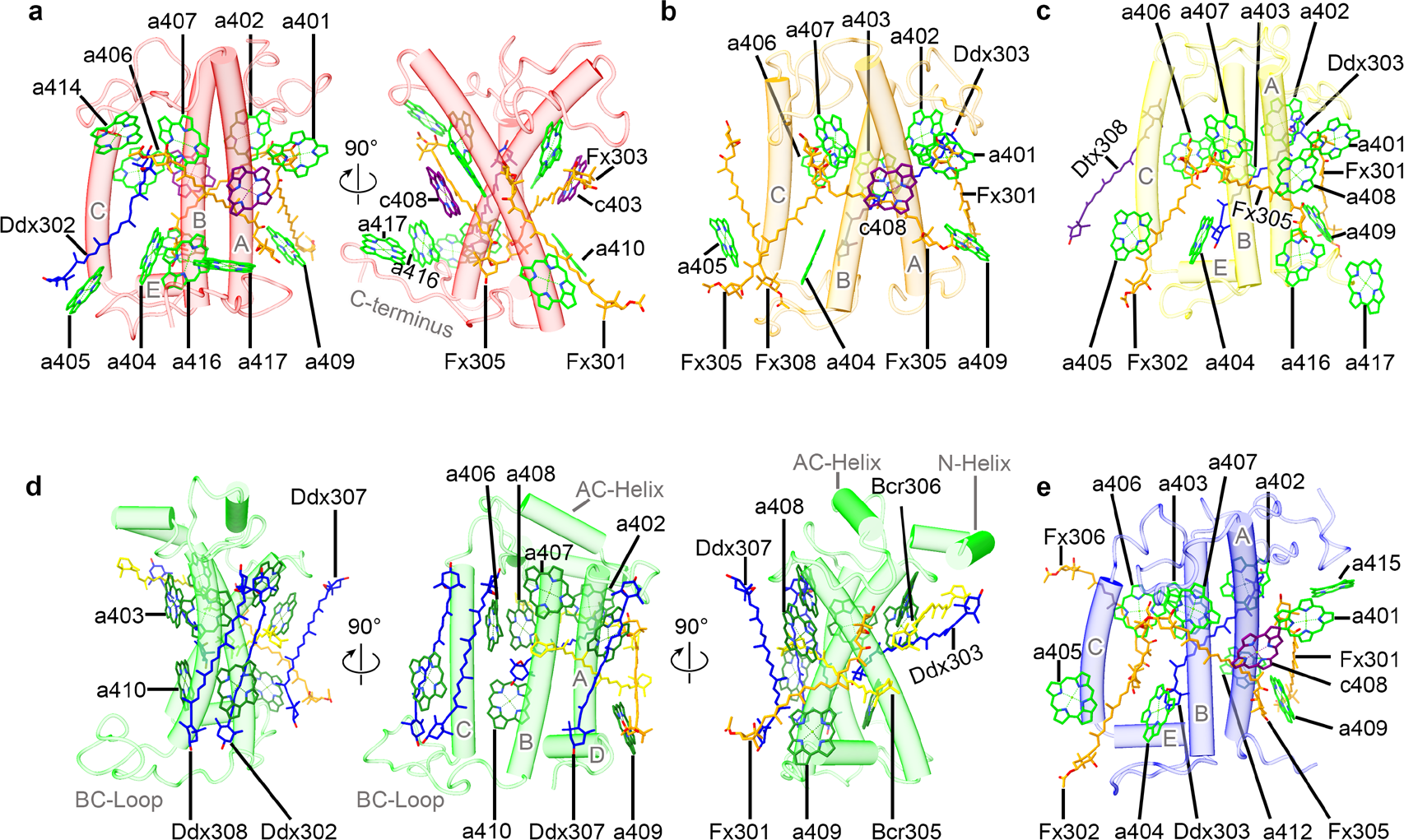
Structures and pigment arrangements of the five FCPIs in PSI-FCPI-L/S. The pigments of Chl *a*, Chl *c*, Fx, Ddx, and Dtx are colored “green,” “purple,” “orange,” “blue,” and “indigo,” respectively, and shown in sticks. Only rings of the Chls are depicted, and their phytol chains are omitted. All pigment molecules are labeled. **a**, Structure and pigment distribution of Tp-Lhcf10. **b**, Structure and pigment distribution of Tp-Lhcq8. **c**, Structure and pigment distribution of Tp-FCP3. **d**, Structure and pigment distribution of Tp-RedCAP **e** Structure and pigment distribution of Tp-Lhcr3

### Structures of the five FCPIs in PSI-FCPI-L/S

According to the sequence phylogenetic analysis (Supplementary Fig. 4), Tp- Lhcf10 belongs to the Lhcf family, Tp-Lhcf10 has a long C-terminus but no Helix-D which is shown as horizontal extension and participates in the interaction with PsaL and Lhcq8 (Fig. 3a and Supplementary Fig. 3a). Tp-Lhcf10 binds 13 Chls and 4 carotenoids (Cars), including 11 Chls *a* (401-402, 404-407, 409-410, 414, 416-417), 2 Chls *c* (403, 408), 3 Fxs (301, 303, 305) and 1 Ddx302, among which 11 Chls (401- 410, 414) are conserved in Cg-Lhcf1^32^ and Tp-Lhcx6_1^33^, the extra Chl *a*416 and Chl *a*417 located between Helix A and long C-terminus (Fig. 3a).

Compared with homologous Cg-Lhcf3 (named by phylogenetic relationships between *C. gracilis* and *T. pseudonana*^31^) in Cg-PSI-24FCPI (Supplementary Fig. 4). Not only the Tp-Lhcf10 site has a big shift at 6.8 Å (Supplementary Fig. 3a) but also has a high rmsd value (RMSD / Cα: Root means square deviation values between two secondary structures of subunits are calculated by Chimera) between Tp-Lhcf10 and Cg-Lhcf3 at 3.75/159 (Supplementary Table 4), Tp-Lhcf10 has 3 conserved transmembrane helices but shorter BE-Loop and so much longer C-terminus, the secondary structures of N-terminus and AC-Loop regions are also different (Supplementary Fig. 5a). In corporations of pigments sites, Tp-Lhcf10 has four more Chls (410, 414, 416, 417) but three fewer Cars (304, 306, 307), the Chl *a*410 and *a*414 in Tp-Lhcf10 occupy the locations of Fx304 and Fx306 in Cg-Lhcf3, respectively (Supplementary Fig. 5a). The extra Chl *a*416 and *a*417 surround with Long C-terminus in Tp-Lhcf10 which occupies the location of Fx307 in Cg-Lhcf3. The shift Chl *a*405 and Car302 in Tp-Lhcf10 was caused by a shorter BE-Loop and the Fx302 in Cg-Lhcf3 was replaced by Ddx302 in Tp-Lhcf10 (Supplementary Fig. 5a). In this structural heterogeneity Lhcf site, Tp-Lhcf10 combines more Chls and Ddx but fewer Fxs, which could facilitate faster energy transfer and quenching and avoid collecting too much blue-green light compared with Cg-Lhcf3.

Tp-Lhcq8 belongs to the Lhcq family (Supplementary Fig. 4), most other Lhcq FCPIs have proven combined in the outer belt^22,31^, this conserved site was found in Cg- PSI-24FCPI and built by Cg25468 in transcriptome data which correspond to Cg- Lhcq12 based on genome^31,36^, Tp-Lhcq8 binds 9 Chls and 5 Cars, including 8 Chls *a* (401-407, 409), 1 Chl *c*408, 4 Fxs (301, 302, 305, 308) and 1 Ddx303 (Fig. 3b and Supplementary Table 2). The rmsd value between Tp-Lhcq8 and Cg-Lhcf12 is 1.64/155 (Supplementary Table 4), Tp-Lhcq8 has a longer C-terminus and their BE-Loop is closer to the lumenal side. All the Chl sites between them are conserved but Tp-Lhcq8 has one more Fx308 for their BE-loop closer to the lumenal side could accommodate the head area of the additional Fx308 (Supplementary Fig. 5b) and the Fx303 in Cg- Lhcq12 was replaced by Ddx303 in Tp-Lhcq8 (Supplementary Fig. 5b).

Tp-FCP3 is homologous with Cg-Lhcr9 (Supplementary Fig. 4a, b), also named Tp-Lhcq10 based on phylogenetic relationships between *C. gracilis* and *T. pseudonana*^31^, among which make up an independent clade (Supplementary Fig. 4c) (Cg-Lhcr9 clade in the previous study), this clade can not be classified between Lhcr, Lhcq and Lhcx, so we named it Tp-FCP3 by its original gene name FCP3 (UniProt ID: B8CEQ3) according to the genome^27^. Tp-FCP3 binds 11 Chls and 5 Cars, including 11 Chls *a* (401-409, 416-417), 3 Fxs (301, 302, 305), 1 Ddx303 and 1 Dtx308 (Fig. 3c and Supplementary Table 2). The rmsd value between Tp-FCP3 and Cg-Lhcr9 is 1.44/161 (Supplementary Table 4), Tp-FCP3 has two more amino acids than Cg-Lhcr9 in the C- terminus thus coordinated with extra Chl *a*417 by Leu194 (Supplementary Fig. 5c), an additional diatoxanthin molecule assigned to Car308 site which is not found in Cg-PSI- FCPI^21,22^ may participate in energy quenching (Fig. 3c and Supplementary Fig. 5c).

Tp-RedCAP belongs to the RedCAP family (Supplementary Fig. 4c) and was built by WWT48828.1 found in the Intron of the genome^27^. The location of Tp-RedCAP is conserved compared with FCPI-1 (fc13194) from *C. gracilis*^21^, RedCAP from *P. purpureum*^18^ and ACPI-8 from *C. placoidea*^20^, both of them have conserved 3 transmembrane helices and amphipathic helix D but special N-Helix and extended BC- Loop compared with LHC proteins. In addition, Tp-RedCAP forms a unique AC-Helix rather than the typical AC-Loop in all LHC proteins (Fig. 3d), the N-Helix and AC- Helix were not built in Cg-FCPI-1 because of their limited local map and transcriptome data^21^. Tp-RedCAP only binds 8 Chls *a* (402-403, 405-410) but 7 Cars(301-303, 305- 307) (Fig. 3d and Supplementary Table 2), the Chl *c*408 may be wrongly assigned in Cg-FCPI-1 limited by the loss of the area of the density map between AC-Helix and Helix A which was found to overlap with the characteristic polar C-17 propionic acids in Chl *c* (Supplementary Fig. 5d), we assign Chl *a* in 408 site in this study. All of its Chl sites assigned by Chl *a* but no Chl *c* could better correspond to the original definition of RedCAP^43,44^ (red lineage chlorophyll *a*/*b*-binding-like protein), all the Chl sites modeled Chl *a* is similar to the RedCAP found in other red-lineage organisms^18,20^, Tp-RedCAP has four more amino acids in the C-terminus than Cg-FCPI-1 thus binds the extra Chl *a*409. More sites of Cars were found in Tp-RedCAP compared with Cg-FCPI-1 based on the better map, including 1 Fx301, 2 Bcrs (305, 306) and 4 Ddxs (302, 303, 307, 308), which indicates that Tp-RedCAP could play an important role in energy quenching (Supplementary Fig. 5d).

Tp-Lhcr3 belongs to the Lhcr family (Supplementary Fig. 4c) which binds 11 Chls and 5 Cars, including 10 Chls *a* (401-407, 409, 412, 415), 1 Chl *c*408, 4 Fxs (301, 302, 305, 306) and 1 Ddx303 (Fig. 3e and Supplementary Table 2). The rmsd value between Tp-Lhcr3 and Cg-Lhcr1 is 1.04/163 (Supplementary Table 4), Tp-Lhcr3 has four more amino acids in the C-terminus than Cg-Lhcr1 which occupies the location of Fx301 in Cg-Lhcf1 thus causes large shifts of Fx301 in Tp-Lhcr3, an extra Fx302 embed into grooves formed by the crossing helices B and C, the His184 in Tp-Lhcr3 replaces Val in Cg-Lhcr1 thus coordinate with extra Chl *a*412 (Supplementary Fig. 5e).

### Structures of the eight FCPIs in PSI-FCPI-Large

Compared with PSI-FCPI-L, PSI-FCPI-S loss 8 Lhcr family FCPIs, which are homologous with Cg-Lhcr2-8 and Cg-Lhcr10 in Cg-PSI-FCPI (Supplementary Fig. 4). The content of PSI-FCPI-S increased apparently in high-light (Fig. 1d) indicating that the 8 Lhcr family FCPIs may respond to high-light conditions and thus detached from the PSI-FCPI supercomplex.

Tp-Lhcr14 binds 13 Chls and 4 Cars, including 12 Chls *a* (Chls 401-407, 409, 410, 412, 415, 421), 1 Chl *c*408, 2 Fxs (302, 305) and 2 Ddxs (303, 306) (Fig. 4a and Supplementary Table 2). The rmsd value between Tp-Lhcr14 and Cg-Lhcr2 is 1.34/170 (Supplementary Table 4), the numbers and sites of Cars are conserved but the locations of Car305 and Car306 were interchanged between Fx and Ddx (Supplementary Fig. 5f). Tp-Lhcr18 binds 9 Chls and 3 Cars, including 8 Chls *a* (402-407, 410, 418), 1 Chl *c*408 and 3 Ddxs (302, 303, 308) (Fig. 4b and Supplementary Table 2). The rmsd value between Tp-Lhcr18 and Cg-Lhcr3 is 1.36/150 (Supplementary Table 4), the His94 in Tp-Lhcr18 replaces Ser in Cg-Lhcr3 thus coordinates with extra Chl *a*418 (Supplementary Fig. 5g). Tp-Lhcr18 loss the Chl *a*415, Fx301, 305, 306 in Cg-Lhcr3 and three Ddxs replace the Fx sites (302, 303, 308) (Supplementary Fig. 5g), which indicates that Tp-Lhcr18 has more capability in energy quenching but less in blue-green light harvesting.

**Fig. 4.**
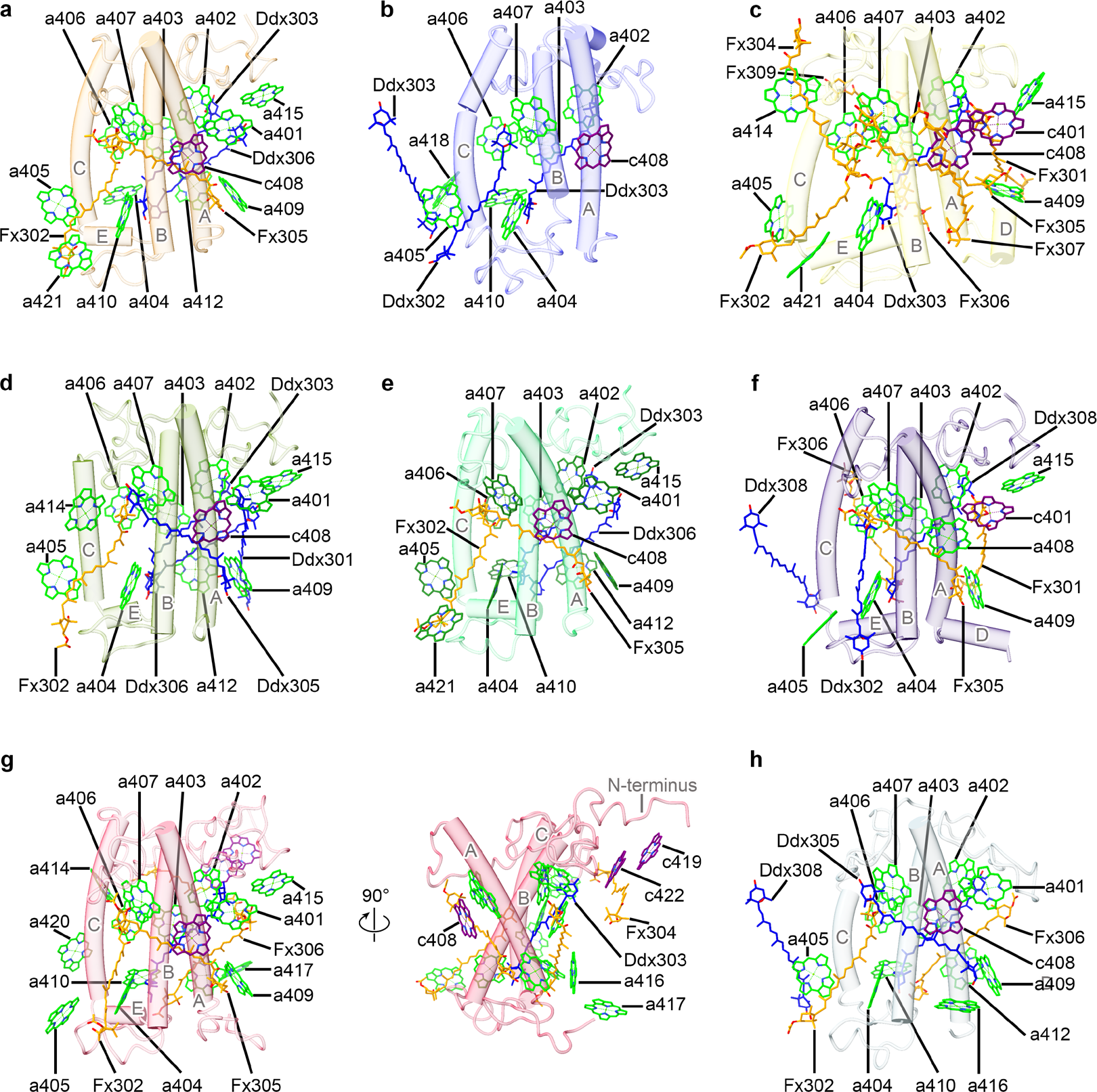
Structures and pigment arrangements of the eight FCPIs in PSI-FCPI-L. The pigments are colored and shown as in Fig. 3. **a**, Structure and pigment distribution of Tp-Lhcr14. **b**, Structure and pigment distribution of Tp-Lhcr18. **c**, Structure and pigment distribution of Tp-Lhcr10. **d**, Structure and pigment distribution of Tp-Lhcr1. **e**, Structure and pigment distribution of Tp-Lhcr4**. f**, Structure and pigment distribution of Tp-Lhcr7**. g**, Structure and pigment distribution of Tp-Lhcr11**. h**, Structure and pigment distribution of Tp-Lhcr20.

Tp-Lhcr10 binds 12 Chls and 8 Cars, including 10 Chls *a* (402-407, 409, 414, 415, 421), 2 Chls *c* (401, 408), 7 Fxs (301, 302, 304-307, 309) and 1 Ddx303 (Fig. 4c and Supplementary Table 2). The rmsd value between Tp-Lhcr10 and Cg-Lhcr4 is 1.23/173 (Supplementary Table 4) which leads to differences in N-terminus and BE-Loop areas, among which coordinate with extra Chl *a*415 and Chl *a*421 in Tp-Lhcr10, respectively (Supplementary Fig. 5h).

Tp-Lhcr1 binds 12 Chls and 5 Cars, including 11 Chls *a* (401-407, 409, 412, 414, 415), 1 Chl *c*408, 1 Fx302 and 4 Ddxs (301, 303, 305, 306) (Fig. 4d and Supplementary Table 2). The rmsd value between Tp-Lhcr1 and Cg-Lhcr5 is 0.97/168 (Supplementary Table 4), the Glu128 in Tp-Lhcr1 replaces Leu in Cg-Lhcr5 thus coordinating with extra Chl *a*414 and the Chl *a*414 site occupy the location of Fx302 in homologous Cg-Lhcr5 (Supplementary Fig. 5i).

Tp-Lhcr4 binds 13 Chls and 4 Cars, including 12 Chls *a* (401-407, 409, 410, 412, 415, 421), 1 Chl *c*408, 2 Fxs (302, 305) and 2 Ddxs (303, 306) (Fig. 4e and Supplementary Table 2). The rmsd value between Tp-Lhcr4 and Cg-Lhcr6 is 1.02/170 (Supplementary Table 4), the Fx305 replaces Ddx305 for the shorter AC-loop in Tp- Lhcr4 could accommodate the head area of the replaced Fx308 (Supplementary Fig. 5j). Tp-Lhcf7 binds 10 Chls and 6 Cars, including 9 Chls *a* (402-409, 415), 1 Chl *c*401, 3 Fxs (301, 305, 306) and 3 Ddxs (302, 303, 308) (Fig. 4f and Supplementary Table 2). The rmsd value between Tp-Lhcr7 and Cg-Lhcr7 is 1.19/182 (Supplementary Table 4), Tp-Lhcr7 loses an Fx307 because of its loose binding compared with Cg-Lhcr7 surrounded by outer belt FCPIs in Cg-PSI-24FCPI (Supplementary Fig. 5k).

Tp-Lhcr11 binds 17 Chls and 5 Cars, including 14 Chls *a* (401-407, 409, 410, 414-417, 420), 3 Chls *c* (408, 419, 422), 4 Fxs (302, 304-306) and 1Ddx303 (Fig. 4g and Supplementary Table 2). The rmsd value between Tp-Lhcr11 and Cg-Lhcr8 is 1.76/208 (Supplementary Table 4). Tp-Lhcr11 has a longer C-terminus thus coordinating with an extra *a*417and the Ser162 in Tp-Lhcr11 replaces Leu in Cg-Lhcr8 thus coordinating with extra Chl *a*420 (Supplementary Fig. 5l). The Chl *a*401 in Tp-Lhcr11 replaces the Chl *c*401 in Tp-Lhcr8 for the Thr208 in Tp-Lhcr11 replaces Arg in Tp-Lhcr8 thus can not coordinate with the polar C-17 propionic acids of Chl *c*401 in Tp-Lhcr8, and the Chl *c* move to 408 site and coordinate with Lys212 in Tp-Lhcr11 (Supplementary Fig. 5l).

Tp-Lhcr20 binds 12 Chls and 5 Cars, including 11 Chls *a* (401-407, 409, 410, 412, 416), 1 Chl *c*408, 2 Fxs (302, 306) and 3 Ddxs (303, 305, 308) (Fig. 4h and Supplementary Table 2). The rmsd value between Tp-Lhcr20 and Cg-Lhcr10 is 1.69/163 (Supplementary Table 4). Tp-Lhcr20 has a shorter C-terminus but the last His201 coordinates extra Chl *a*416 which occupies the location of four more amino acids in Cg-Lhcr10, the His194 in Tp-Lhcr20 replaces Thr in Cg-Lhcr10 thus coordinate with extra Chl *a*412 which occupy the site of Fx302 in Cg-Lhcr10 (Supplementary Fig. 5m).

### Organization of FCPIs and their interactions with the PSI core

In this study, we analyzed the interaction mechanisms between antennas and between antennas and the core using the VDW method^33^ (see methods for details), two weak interactions were found in Tp-Lhcr1 to Tp-Lhcr10 and Tp-Lhcr20 to Tp-Lhcf10 (Fig. 5a, b) thus formed three thus resulting in three interacted regions with the PSI core. Compared to the antenna belt definition from a recent study^23^, we optimized the definitions of belt 1 and belt 3 in PSI-FCPI through the structures of PSI-FCPI-L and PSI-FCPI-S in this study (Fig. 5b and Supplementary Fig. 6). Five interacted FCPIs in Tp-PSI-FCPI-S formed unique belt 3 (Fig. 2b and Supplementary Fig. 6b). The Tp- PsaR subunit is conserved with rhodophytes and cryptophytes compared to primitive rhodophytes^18,20,23^ (Supplementary Fig. 6a). Therefore, these organisms could bind belt 2 (including three antennas). Tp-Lhcr11 occupies the position of PsaK (Supplementary Fig. 3b), connecting Tp-Lhcr7 and strongly interacting with Tp-Lhcr20, resulting in a continuous belt 1 composed of five FCPIs (Supplementary Fig. 6b). The belt 1 and belt 2 interacted with the PSI core mainly concentrated in the stromal side because of the bigger gap in the lumenal side (Fig. 5b).

**Fig. 5.**
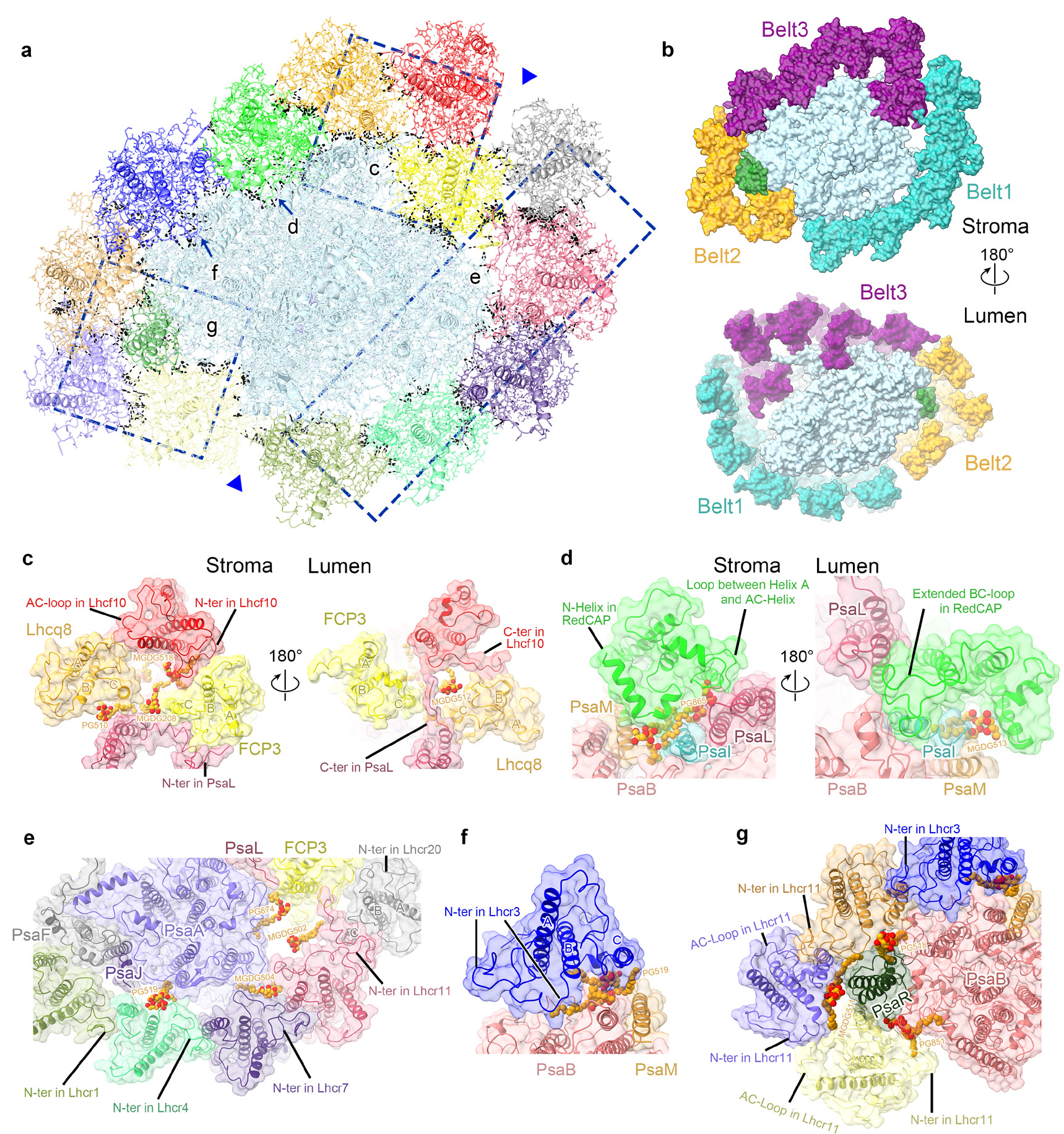
Interactions among different FCPIs and FCPIs to PSI core in the PSI-FCPI supercomplex from *T. pseudonana*. The interaction subunits were shown in a transparency surface mode and all lipids were shown as orange spheres. O atoms, red. **a**, Van der Waals forces analysis in adjacent FCPIs and FCPIs to PSI core. The black dashed lines represent interactions and the blue triangles point to the gap caused by weak interactions. **b**, Three FCPI belts interacted with the PSI core. The belt 1 and belt 2 interacted mainly in the stromal side. **c**, Interactions between Tp-Lhcf10, Tp-Lhcq8, Tp-FCP3 and PSI core. **d**, Interactions between Tp-RedCAP to PSI core. **e**, Interactions between the belt 1 to PSI core. **f**, Interactions between Tp-Lhcr3 to PSI core. **g** Interactions between the belt 2 to PSI core.

The interactions between Tp-Lhcf10 and PSI core are different from Cg-Lhcf3 and PSI core (Supplementary Fig. 3a). The N-terminus and longer C-terminus in Tp-Lhcf10 form interactions with Tp-FCP3, Tp-Lhcq8 and longer Tp-PsaL. In addition, MGDG518 of Tp-Lhcf10 and MGDG512 of Tp-Lhcq8 also strengthen the interactions. Tp-Lhcq8 interacts with the PSI core mainly through its Helix C connected to Tp-PsaL with the help of two lipids in the stroma, PG510 of Tp-Lhcq8 and MGDG208 of PsaL, respectively (Fig. 5c).

Tp-FCP3 interacts with the PSI core mainly through its AC-Loop connected to the N-terminus of Tp-PsaL with the help of MGDG208 (Fig. 5c). Tp-RedCAP interacts with the PSI core through N-Helix and Loop area connect to PsaB, PsaI, PsaL and PsaM with the help of PG865 in the stromal side, the extended BC-Loop connects to PsaB, PsaI and PsaL in the lumenal side, MGDG513 of Tp-RedCAP strengthen the interactions (Fig. 5d).

Tp-FCP3 covers the location of PsaO and occupies more space thus breaking a gap between Tp-Lhcf10 (belt 3) and Tp-Lhcr20 (belt 1). The belt 1 connects to the PsaA side mainly through the extremely long N-terminus of Tp-Lhcr11 interacts with Tp- FCP3 and PsaA. Four lipids strengthen the interactions between belt 1 and the PSI core in the stromal side (Fig. 5e). Tp-FCP3 plays an important role in binding belt 1 and belt 3.

Tp-Lhcr3 in belt 3 connects to the PsaB and PsaM with the help of PG519 (Fig. 5f). In addition, belt 2 connects to the PsaB side mainly through the PsaR and Tp-Lhcr3 subunit, Tp-Lhcr14 and Tp-Lhcr10 interact with PSI core through their N-terminus, three lipids strengthen the interactions between belt 2, PsaR and PSI core (Fig. 5g).

### Energy transfer pathways

The outer FCPI belt in the Cg-PSI-24FCPI supercomplex is mainly encoded by lhcq genes^31,36^ (Supplementary Fig. 6b), 11 outer FCPIs (including 9 Lhcqs) of 24 FCPIs contribute 54/102 Fxs (Supplementary Fig. 7a), thus participating in harvesting blue-green light which could enhance the capability in light-harvesting at low light. In this study, the outer FCPI belt was lost in high light conditions and the ratio of PSI-FCPI-S/PSI-FCPI/L increased apparently (Fig. 1d). The belt 1 and belt 2 detached from the PSI-FCPI complex could avoid photodamage caused by FCPIs in high light.

The pigments of the Tp-PSI core are conserved compared with the Cg-PSI core, with a small difference in losing Bcr845 of Tp-PsaB (Fig. 6a, b), both of them have 94 Chls *a*. In addition, most Chls *c* in FCPIs are distributed in the 408 site (Supplementary Table 5) which surrounds the edge of the Tp-PSI-FCPI supercomplex (Fig. 6a). Tp- FCPIs have more Chls *a* compared with the homologous 13 FCPIs in Cg-PSI-24FCPI (Supplementary Fig. 7), some of them occupy the sites of Fxs, and some Ddxs also replace Fxs in several carotenoid sites (Fig. 6b). The additional Chls *a* coordinated with mutant amino acids and thus form different pigment networks due to the different diatom species (Supplementary Fig. 5a, c-e, g-i, l, m).

**Fig. 6.**
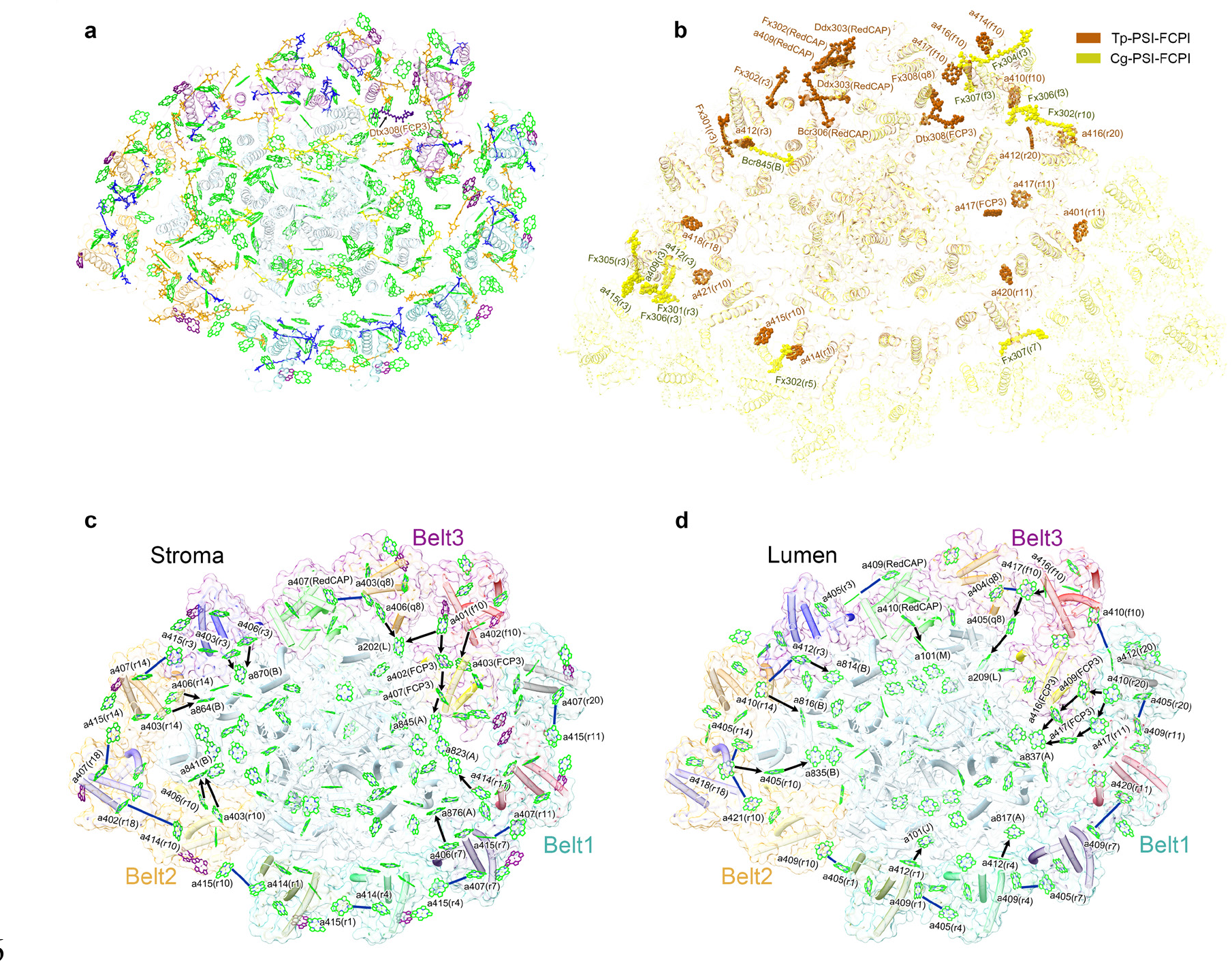
Pigment arrangement and possible EET pathways in the Tp-PSI–FCPI-L supercomplex from *T. pseudonana*. The structure is viewed from the stromal side. The colors of the FCPI subunits and pigments are the same as in Fig. 2 and Fig. 3. The EET pathways from FCPIs to PSI core were indicated by black arrows and the energy transfer pathways between FCPIs were indicated by blue lines. **a**, Distribution and arrangement of all pigments in the Tp-PSI–FCPI-L supercomplex. **b**, The extra and missing pigment sites compared with Cg-PSI-FCPI were labeled in spheres. **c**, EET pathways in the Tp-PSI–FCPI-L in the stromal side. **d**, EET pathways in the Tp-PSI– FCPI-L in the lumenal side.

The excited energy transfer (EET) pathways were analyzed by the Föster resonance energy transfer (FRET) method^12,45^, and we demonstrate the EET pathways from FCPs to the PSI core at the stromal and lumenal sides, respectively (Fig. 6c, d).

The efficient EET pathways from FCPIs to PSI core are demonstrated by the pathways that lifetime<10 ps and the energy transfer pathways between FCPIs could demonstrate with the pathways that lifetime<40 ps (Supplementary Fig. 8).

The EET pathways from belt 1 to the PSI core include Chl *a*412_Lhcr1_→Chl *a*101_PsaJ_, Chl *a*412_Lhcr4_→Chl *a*817_PsaA_, Chl *a*406_Lhcr7_→Chl *a*876_PsaA_, Chl *a*414_Lhcr11_→Chl *a*823_PsaA_ (Fig. 6d), the Tp-Lhcr20 far from the PSI core and transfer their energy mainly through Chl *a*410_Lhcr20_→Chl *a*409_FCP3_→Chl *a*416_FCP3_→Chl *a*837_PsaA_ and rely on two additional Chls *a*417 form the unique Chl *a*410_Lhcr20_→Chl *a*417_Lhcr11_→Chl *a*417_FCP3_→ Chl *a*837_PsaA_ pathway (Fig. 6d). Energy transfer pathways between FCPIs in belt 1 including Chl *a*415_Lhcr1_ and Chl *a*407_Lhcr4_, Chl *a*415_Lhcr4_ and Chl *a*407_Lhcr7_, Chl *a*415_Lhcr7_ and Chl *a*407_Lhcr11_, Chl *a*415_Lhcr11_ and Chl *a*407_Lhcr20_ at the stromal side (Fig. 6c), Chl *a*409_Lhcr1_ and Chl *a*405_Lhcr4_, Chl *a*409_Lhcr4_ and Chl *a*407_Lhcr7_, Chl *a*409_Lhcr7_ and Chl *a*420_Lhcr11_, Chl *a*409_Lhcr11_ and Chl *a*405_Lhcr20_ at the lumenal side (Fig. 6d).

The EET pathways from belt 2 to the PSI core include coupled Chls *a*403- *a*406_Lhcr14_→Chl *a*864_PsaB_, Chls *a*403-*a*406_Lhcr10_→Chl *a*841_PsaB_ at the stromal side and Chl *a*410_Lhcr14_→Chl *a*816_PsaB_, Chl *a*405_Lhcr10_→Chl *a*835_PsaB_ at the lumenal side (Fig. 6c). Tp-Lhcr18 lost three Chls *a*409, *a*412 and *a*415 which could accept the energy from outer Lhcqs (Fig. 6b), Tp-Lhcr18 transfers their energy to PSI core mainly through additional Chl *a*418 to Chl *a*405_Lhcr10_ at the lumenal side (Fig. 6d). Energy transfer pathways between FCPIs in belt 2 including Chl *a*415_Lhcr14_ and Chl *a*407_Lhcr18_, Chl *a*402_Lhcr18_ and Chl *a*414_Lhcr10_ at the stromal side, Chl *a*405_Lhcr14_, Chl *a*418_Lhcr18_ and Chl *a*421_Lhcr10_ at the lumenal side (Fig. 6c, d).

The EET pathways from belt 3 to the PSI core include coupled Chls *a*403-*a*406_Lhcr3_ →Chl *a*870_PsaB_, Ch *a*406_Lhcq8_→Chl *a*202_PsaL_ at the stromal side and Chl *a*412_Lhcr3_→Chl *a*814_PsaB_, Chl *a*410_RedCAP_→Chl *a*101_PsaM_, Chl *a*405_Lhcq8_→Chl *a*209_PsaL_ at the lumenal side (Fig. 6c, d). Tp-Lhcf10 transfer their energy through Chl *a*404_Lhcf10_→Chl *a*202_PsaL_, Chls *a*401, *a*402_Lhcf10_→Chl *a*402, *a*403_FCP3_→Chl *a*407_FCP3_→Chl *a*845_PsaA_ at the stromal side and Chl *a*416, *a*417_Lhcf10_→Chl *a*405_Lhcq8_→Chl *a*209_PsaL_ at the lumenal side.

Energy transfer pathways between FCPIs in belt 3 including Chl *a*407_RedCAP_ and Chl *a*403_Lhcq8_ at the stromal side, Chl *a*405_Lhcr3_ and Chl *a*409_RedCAP_, Chl *a*404_Lhcq8_ and Chl *a*417_Lhcf10_ at the lumenal side (Fig. 6c, d).

The Energy transfer pathways between belt 1 and belt 2 include Chl *a*414_Lhcr1_ and Chl *a*415_Lhcr10_ at the stromal side, Chl *a*405_Lhcr1_ and Chl *a*409_Lhcr10_ at the lumenal side. The Energy transfer pathways between belt 2 and belt 3 include Chl *a*407_Lhcr14_ and Chl *a*415_Lhcr3_ at the stromal side, Chl *a*410Lhcr14 and Chl *a*412_Lhcr3_ at the lumenal side. The Energy transfer pathways between belt 1 and belt 3 through Chl *a*412_Lhcr20_ and Chl *a*410_Lhcf10_ at the lumenal side (Fig. 6c, d).

Compared with homologous Cg-PSI-13FCPI (Supplementary Fig. 7), Tp-PSI- FCPI has more Chls *a* and Ddx/Dtx but fewer Fxs, 10 additional Chls *a* enhance the capability of energy balance between peripheral FCPIs including Chl *a*414_Lhcr1_, Chl *a*420_Lhcr11_, Chl *a*412_Lhcr20_, Chl *a*418_Lhcr18_, Chl *a*421_Lhcr10_, Chl *a*415_Lhcr10_, Chl *a*410_Lhcf10_, Chl *a*417_Lhcf10_, Chl *a*409_RedCAP_ and Chl *a*412_Lhcr3_ (Fig. 6c, d). Two additional Chl *a*417_Lhcr11_ and Chl *a*417_FCP3_ involve energy transfer from belt 1 to the PSI core (Fig. 6d). The RedCAP is rich in Bcrs and Ddxs which may play an important role in quenching excess energy (Fig. 3d), and the Dtx308 found in FCP3 is located between stable binding belt 3 and PSI core (Fig. 6a, b), we speculate that this Dtx may serve as an efficient energy-quenching site (Fig. 6b).

### Potential assembly mechanisms and light-adaptive strategies in diatom PSI-FCPI

Diatoms survive in oceanic environments with fluctuating light conditions that need to strengthen light-harvesting capability in low light but avoid photodamage caused by excess light energy in high light^46^. The previous study proved that the PSI core of diatom *C. gracilis* could bind 16-24 FCPIs to harvest light energy underwater in low light^21,22^. In this study, two smaller PSI-FCPI-L/S complexes from *T. pseudonana* were resolved by Cryo-EM (Fig. 2 and Supplementary Fig. 1), both of them lost outer Belts composed of Lhcqs, and the ratio of PSI-FCPI-S/PSI-FCPI-L increased under high light conditions (Fig. 1d).

belt 1 and belt 2 (8 Lhcr FCPIs) could detach from PSI-FCPI-L and form unique PSI-FCPI-S, indicating that the diatom PSI core could avoid photodamage through reduced peripheral FCPIs under high light conditions. The outer Lhcq belt was not found in this study, but the homologous *lhcq* genes that exist in *T. pseudonana* prove that the PSI-FCPI also could assemble the outer Lhcq FCPIs (Supplementary Fig. 4c), such as the potential PSI-18FCPI in the previous study^35^ or even huge PSI-24FCPI^21^.

The Tp-RedCAP and Tp-FCP3 rich in Ddx/Dtx (Fig. 3c, d) may participate in energy quenching from Tp-Lhcf10 and Tp-Lhcq8, this may be one of the reasons why belt 3 could bind with the PSI core stably. In addition, Tp-Lhcr3 is located in a conserved antenna site in the red lineage PSI supercomplexes different from the flexible RedCAP site^19^ and others (Supplementary Fig. 6).

We further analyzed the reasons why Tp-Lhcr3 could combine with PSI core stably, one conserved Lhcr protein named HLR1^47^ from *Nannochloropsis oceanica* was found in high homology with Tp-Lhcr3 or homologous Cg-Lhcr1 in diatom (Supplementary Fig. 9). Both *Eustigmatophyceae* (including Nannochloropsis), *Phaeophyceae* (brown algae) and *Bacillariophyceae* (diatoms) belong to stramenopiles (Heterokontophyta) in red lineage organisms^47,48^. HLR1 has been proven to play an important role in the dynamic balance between photoprotection and efficient light harvesting in photosynthesis, the *hlr1* mutant could avoid photodamage in high light but their mature PSI complex lost more antennas^47^. Therefore, we speculate that conserved homologous Tp-Lhcr3 antenna is also involved in starting to assemble peripheral FCPIs even if the combination with the PSI core in high light will cause photodamage. The TpLhcr3 combines belt 2 to the PsaB side with the help of the PsaR subunit (Fig. 5g). On the other hand, the Tp-FCP3 plays an important role in assembling belt 1 and TpLhcf10 belt 3 to the PsaA side by interacting with the N-terminus of Tp-Lhcr11 and Tp-Lhcf10 (Fig. 5c, e).

The two conserved large subunits PsaA and PsaB, among which demonstrate a pseudosymmetric structure that evolved from an ancient homodimeric assembly^1,13^. In the green lineage PSI supercomplex, the PsaK and PsaG combine the LHCI belts to PsaA and PsaB sides, respectively^49^. The PsaO and PsaH combine the LHCII trimer to the PSI core^14,15^. In diatom, the FCPI sites of TpLhcr3 and TpFCP3 in belt 3 participate in combining FCPIs in belt 1 and belt 2 to PsaA and PsaB sides with the help of unique PsaR in red lineage organisms (Fig. 5). In addition, three FCPI sites of Tp-RedCAP, Tp-Lhcq8 and Tp-Lhcf10 also could be detached from the PSI-16FCPI supercomplex^22^, indicating that the two FCPI sites of TpLhcr3 and TpFCP3 are most likely to serve as the original assemble sites in the PsaA and PsaB sides, respectively.

Based on the summarized points and two structures of PSI-FCPI-L/S resolved in high light, the assembly mechanisms and light-adaptive strategies in the diatom PSI-FCPI supercomplex may follow the following steps (Fig. 7). First, the Tp-FCP3 and Tp-Lhcr3 combine to PsaA and PsaB sides, respectively. Then, the five FCPIs in belt 3 were assembled at the PsaL side. belt 1 and belt 2 combined to PsaA and PsaB sides with the help of Tp-FCP3 and Tp-Lhcr3, Tp-Lhcr11 and PsaR that strengthen the capability in light-harvesting. A large number of Lhcq FCPIs in the outer belt surround the PSI-FCPI-L to further strengthen the capability in light-harvesting under low light. The Lhcq FCPIs and 8 Lhcr FCPIs could detached from the PSI-FCPI supercomplex under high light conditions to avoid photodamage (Fig. 7). The status of PSI-FCPI-S could again efficiently combine more peripheral antennas from high light to low light. The assembly mechanisms and light-adaptive strategies in the PSI-FCPI supercomplex could promote diatom survival in fluctuating oceanic environments.

**Fig. 7.**
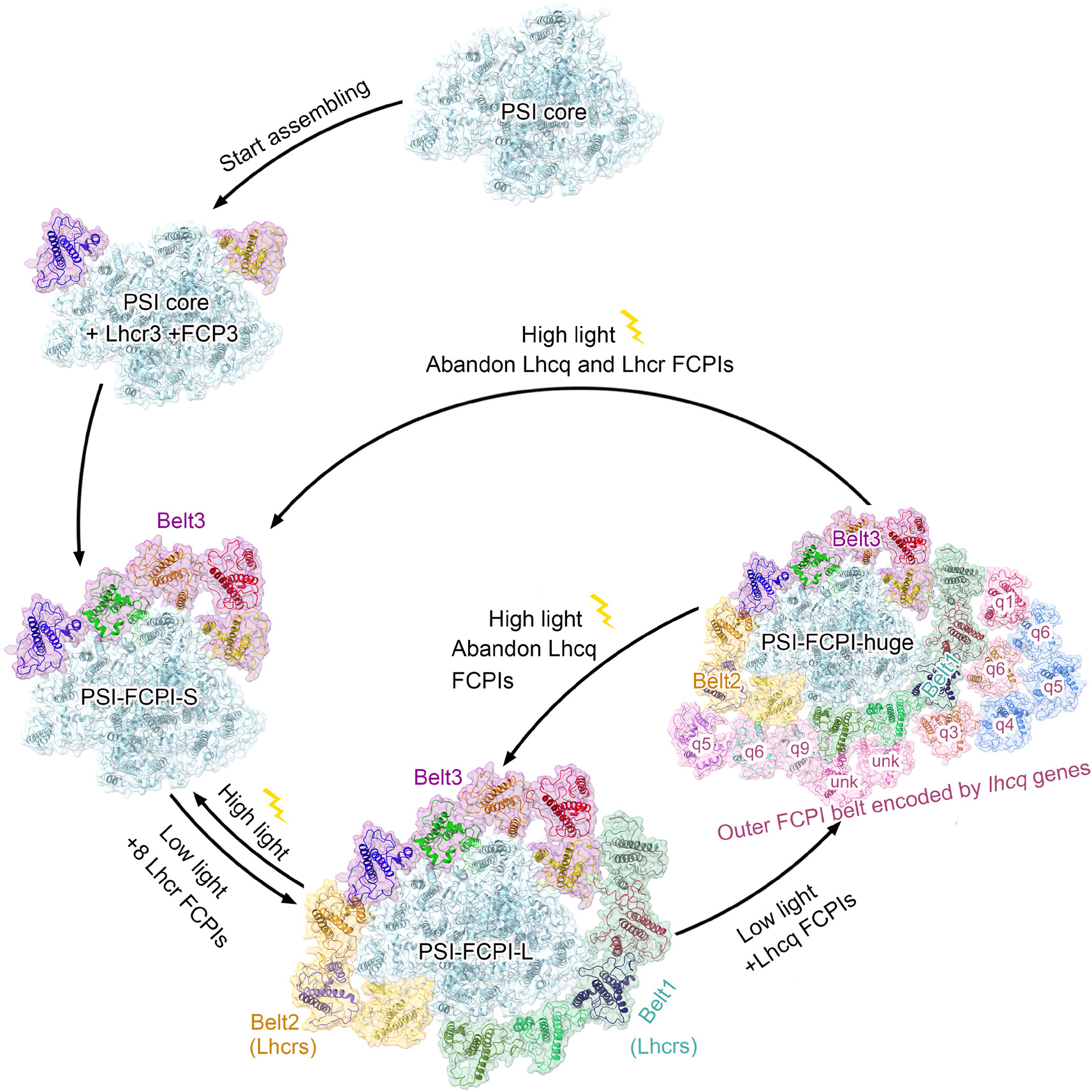
Potential light-adaptive strategies in diatom PSI-FCPI. The Tp-FCP3 and Tp-Lhcr3 combine to PsaA and PsaB sides play important roles in connecting peripheral FCPIs, stable FCPIs in belt 3 could be assembled firstly, the eight Lhcr FCPIs and the outer FCPIs mainly composed of Lhcqs could combined flexibly and influenced under different light conditions. PSI-FCPI-huge was modeled by PDB: 6LY5

## Discussion

The peripheral antennas of the PSI supercomplex are important for light harvesting, transfer and dissipation^23,50^. Diatoms combine more lhc families in PSI-FCPI than green lineage PSI supercomplex^28^, including Lhcf, Lhcr, Lhcq, unique FCP3 and RedCAP (Supplementary Fig. 4). This indicates that the PSI diatom could achieve more functions by combining different types of antennas.

The previous studies identified the FCPI subunits^51^ in Tp-PSI-FCPI and solved the structure of Tp-PSI-FCPI by Cryo-ET and EM in 17 Å^35^, among which speculated that the PSI-FCPI demonstrate convergent evolution different from structural heterogeneity in PSII-FCPII. Most previously identified FCPIs^51^ are well matched in this study based on the 2.78 Å resolution except speculated Lhcx6_1^51^ which was proven combined with PSII-FCPII^33^. The final Cryo-EM structures of Tp-PSI-FCPI-L/S in this study are similar to the two smaller 2D classification results of PSI-FCPI^35^, but the previously identified Cryo-ET model of Tp-PSI-18FCPI was not found under high light conditions. Recently, the relative abundances of FCPI composition were found prominent change in high light effect in Tp-PSI-FCPI^36^. We analyzed the molecular weight of PSI-FCPI supercomplex purified under different light conditions by gel filtration chromatography, more PSI-FCPI-L loses 8 Lhcr FCPIs thus forming increased PSI-FCPI-S under high light conditions (Fig. 1d), among which could correspond to the increase of Tp-FCP3, Tp-Lhcr3, Tp-Lhcf10 and decrease of Tp-Lhcr1, Tp-Lhcr7 and TpLhcr10 from low light to high light in the recent study^36^. The structures of PSI-FCPI-L/S in the present study can not prove the change of Tp-Lhcr4, Tp-Lhcr14 and Tp-Lhcq8 (FCP10)^36^. Three Tp-Lhcr18, Tp-Lhcr20 and Tp-RedCAP were not annotated in the UniProt database thus were not analyzed their changes^36^, but a homologous LHL1 protein of RedCAP exhibited no significant signs of sudden changes in light conditions in another pennatae diatom *Phaeodactylum tricornutum*^43^. In addition, the Dtx308 was modeled in the speculated Dtx binding candidate Tp-FCP3^36^, located between belt 3 and PSI core and may serve as an efficient energy-quenching site.

The typical phosphorylation-related state transitions was proposed absent in diatoms^28,39^. Compared with the PSI-LHCI-LHCII supercomplex in green lineage organisms^14,15^, the Tp-Lhcf10 occupies the location of LHCII with the help of Tp- Lhcq8 and Tp-FCP3 that occupy the locations of PsaH and PsaO, we do not find any phosphorylated density maps around the three FCPIs area, the Tp-Lhcf10 in belt 3 tend to stable bind with PSI complex and regulate the excited energy from FCPI to the core. However, the three FCPI sites of Cg-Lhcf3 (Tp-Lhcf10), Cg-Lhcq12 (Tp-Lhcq8) and RedCAP were lost in Cg-PSI-16FCPI^22^ supercomplex compared with Cg-PSI- 24FCPI^21^ and Tp-PSI-FCPI, except possible different growth conditions including CO_2_ concentrations and temperatures^40^, another reason may be associated with the treatment of TritonX-100 to thylakoid to separate the PSII and PSI^41,42^, the outer Lhcf site in PSI complex whether associated with PSII do not have enough evidence currently, but this Lhcf site exhibit unique structural heterogeneity (Fig. 3a and Supplementary Figs. 3a and 5a) compared with other conserved Lhcr FCPIs which may correspond to different structural heterogeneity in the PSII-FCPII supercomplex^33,35^.

In this study, the structures of the Tp-PSI-FCPI supercomplex exhibit convergent evolution compared with Cg-PSI-FCPI (Fig. 2 and Supplementary Fig. 4). However, the pigment content and ratio in different diatom PSI-FCPI have differences (Fig. 6b and Supplementary Fig. 7), some mutant amino acids in Tp-PSI-FCPI coordinate with additional Chls *a* (Supplementary Fig. 5a, c-e, g-i, l, m), and 5 additional Chls *a* replace the Fxs, and 10 additional Chls *a* produce extra energy transfer pathways between peripheral FCPIs (Fig. 6c, d). On the other hand, the pigment contents of Ddx and Dtx in Tp-PSI-FCPI are higher than Cg-PSI-FCPI but fewer Fxs (Fig. 6b and Supplementary Fig. 7). We tend to speculate the different pigment contents and ratios were caused by different diatom species, similar phenomena were also found in the PSII-FCPII supercomplex between the two diatoms that more Chls *a* and Ddx/Dtx but less Fxs^33^, the photosynthetic apparatus of *T.pseudonana* may need more pathways for energy quenching and energy balance are needed compared with *C. gracilis*. The Tp- RedCAP and Tp-FCP3 are rich in Bcrs and Ddxs/Dtx (Fig. 3c, d), which could participate in quenching excess energy from FCPIs in stable belt 3, the FCPIs in belt 1 and belt 2 could flexibly combine with the PSI-FCPI under different light conditions (Figs. 1d and 2), among which form the potential unique light light-adaptive strategies in diatom PSI-FCPI (Fig. 7). Further studies could pay attention to the content of RedCAP and the content of N-terminal protein-degrading enzymes in the chloroplast stroma under different light conditions to clarify more light-adaptive mechanisms in centric diatoms.

In conclusion, two structures of PSI-FCPI from *T. pseudonana* with alterable peripheral 8 or 13 FCPIs in this study exhibit the specific arrangement and pigment networks, as well as the convergent evolution of PSI-FCPI in diatoms. 8 Lhcr FCPIs could detach from the PSI-FCPI supercomplex to avoid potential photodamage under high light, 5 FCPIs in considerable diversity could stably combine with the PSI core and quench excess energy through unique RedCAP and FCP3 rich in Ddx/Dtx, and the locations and potential assemble functions of Lhcr3 and FCP3 promote PSI-FCPI efficiently regulate the light-harvesting capability under different light conditions through binding different numbers of FCPIs, among which may be part of the reasons that diatoms become the flourish phytoplankton species in fluctuating oceanic environment.

## Methods

### Cultivation of diatoms under different light conditions

*T. pseudonana* cells (CCMP1335, Guangyu Biological Technology Co. Ltd. Shanghai, China) are cultured in the F/2 medium^52^ and bubbled with 3% CO_2_ and air under stirring at 135 rpm, The cells in low light were cultivated constantly at the light intensity of 30 μmol photons m^−2^ s^−1^ and collected when the concentration reached 6 × 10^6^ cells/liter. The cells grown in high light were cultivated under actual light conditions of 300 μmol photons m^−2^ s^−1^ and collected when the concentration reached 1.2 × 10^6^ cells/liter..

### Purification of PSI-FCPI

Diatom cells were collected by centrifugation at 6,000 g for 15 min and resuspended in an ice-cold buffer of 50 mM 2-morpholinoethanesulfonic acid (Mes)– NaOH (pH 6.5), 10 mM MgCl_2_ and 5 mM CaCl_2_. Cells were disrupted by French Press (3 cycles at 35 MPa), and the thylakoid membranes from diatom cells under low/high light were collected according to the previous method^21,29,33^, thylakoid was resuspended in a buffer of 50 mM Mes-NaOH (pH 6.5), 10 mM NaCl, 5 mM CaCl_2_ (MNC). The thylakoid membranes were solubilized with 1% (w/v) N-dodecyl-α-D-maltoside (α- DDM) (Anatrace, Maumee, OH) for 30 min on ice and centrifuged at 40,000g for 10 min to remove the unsolubilized part. The supernatant was loaded on the linear sucrose density gradient (0.1 to 1.0 M sucrose) in the buffer MNC containing 0.03% α-DDM and centrifuged at 230,000g for 17 hours. Four major bands were obtained after centrifugation, from up to bottom are FCPs, PSI/PSII core, PSI-FCPI and PSII-FCPII. The PSI-FCPI was collected and used for SDS-polyacrylamide gel electrophoresis (PAGE) and pigment analysis.

Further purification by gel filtration chromatography (Cytiva, Superose 6 Increase 10/300 GL) in the buffer of 50 mM Mes- NaOH (pH 6.5), 110 mM NaCl, and 5 mM CaCl_2_ supplemented with 0.03% α-DDM. The mixed elutions (13 ml-15 mL) of Tp- PSI-FCPI-HL were collected and concentrated through ultrafiltration centrifuge tube (molecular weight cutoff: 100 kDa; AMICON, Merck Millipore) to the final concentration of 2 mg Chl/ml. All the procedures were operated under a green light at 4°C or on ice.

### Characterization of PSI-FCPI

The peptides of FCPIs were analyzed by SDS-PAGE by the gel containing 14% polyacrylamide and 7.5 M urea^53^. The gel was stained by Coomassie brilliant blue R- 250 (Sigma-Aldrich).

The room temperature absorption spectra were measured through the ultraviolet-vis spectrophotometer (Shimadzu, Japan), and the fluorescence emission and excitation spectra were measured at 77 K in the fluorescence spectrophotometer (F-7000, Hitachi, Japan) at a Chl concentration of 40 μg Chl/ml.

### Pigment analysis

The pigment compositions of PSI-FCPI supercomplex in low light and high light were analyzed by High-performance liquid chromatography (HPLC) as previously^21,54^. The samples collected from the sucrose density gradient were mixed with 90% (v/v) acetone for 30 minutes at 4°C to extract the pigments^21^. After centrifugation at 12,000g for 15 min, the supernatant was loaded into the C-18 reversed-phase column (5 μm, 100 Å, 250 mm by 4.6 mm, Grace, USA). Pigments were identified based on their absorption spectra and elution times. The HPLC results indicated that the PSI-FCPI supercomplex contained Chl *c*, Fx, Ddx, Dtx, Chl *a*, and Bcr.

### Cryo-EM data processing

The PSI-FCPI supercomplex purified from cells grown under low light and high light were applied to the CryoMatrix Amorphous alloy film R1.2/1.3, 300 mesh grid at the Chl concentration of 2 mg/ml and subsequently vitrified by Vitrobot under 4°C and 100% humidity. The blot time was 3 s with a blot force of -1.

Cryo-EM images were collected with a Titan Krios microscope operated at 300 kV equipped with a Gatan Quantum energy filter, at a slit width of 20 eV, a spherical aberration corrector, a K3 camera (Gatan) operated at a superresolution mode, with × 64,000 magnification. Each movie comprises 32 frames with a total dose of ∼60 e^−^/Å^2^ and a dose rate of 20 e^−^/pixel per s. Data acquisition was performed using the EPU software (Thermo Fisher Scientific) with a defocus range of −1.0 to −2.0 μm. The final images were binned, which resulted in a pixel size of 1.1 Å for further data processing. A total of 5,011 movies from PSI-FCPI under high light were collected and processed with cryoSPARC^55^, resulting in 528,815 particles which were boxed using crYOLO^56^. Particles were extracted from micrographs, and two-dimensional (2D) classification was performed to discard bad particles. After several rounds of 2D classification, 225,477 particles were selected for 3D classification in cryoSPARC^55^. 129,745 particles and 53,013 particles were finally used to construct the structural model of Tp-PSI-FCPI-L and Tp-PSI-FCPI-S under high light at 2.78 Å and 3.20 Å based on the gold-standard FSC with a cutoff value of 0.143 using local refinement in cryoSPARC^55^.

### Model building and refinement

The models of PSI-FCPI-L and PSI-FCPI-S were built based on homologous PSI-FCPI from *C. gracilis* (PDB: 6LY5). Based on the genome of *T.pseudonana* and the phylogenetic relationship between *C.gracilis* and *T.pseudonana*, 13 sequences of Tp- FCPIs were obtained, the structure Tp-PSI core and 13 FCPIs were built by SWISS- MODEL^57^ from Cg-PSI-FCPI and fitted into the 2.78 Å cryo-EM map with UCSF Chimera^58^. Then, all the subunits were checked manually and identified with COOT^37^, such as the Longer C-terminus in Tp-PsaL and Tp-Lhcf10, the unique N-Helix and AC-Helix in RedCAP can be well built based on the 2.78 Å cryo-EM map.

In the process of pigment model building according to the previous study^33,54^, all Chls *c* were built in Chl *c*1 limited by resolution, and the unique polar C-17 propionic acids of Chl *c* could coordinated with closed alkaline amino acids (Supplementary Table 5). All the phytol chains of Chls *a* were modeled according to the actual density. As for the model building of carotenoids, the discrimination of Fx molecules is based on the longer density of the head group in carotenoids (Supplementary Fig. 2c). The discrimination between Ddx and Dtx based on the previous study, Ddxs have a larger thickness in the lower ionon ring caused by the bulge of O atom (epoxy group), the two molecules are similar, and we determined from HPLC that Dtx is present. All the density maps for Ddx/Dtx molecules were checked, and Dtx308 in FCP3 most probably fits with the density map of Dtx obtained in this study (Supplementary Fig. 2c).

The model of PSI-FCPI-S was built by PSI-FCPI-L in this study, we deleted eight peripheral FCPIs and fitted them into the 3.20 Å cryo-EM map with UCSF Chimera. Then, we manually refined all the subunits and ligands in COOT as above.

Finally, the PSI-FCPI-L and PSI-FCPI-S models were refined using Phenix^38^, and the statistics for Cryo-EM data and structure refinement are summarized in Supplementary Table 1. The structures in this paper were displayed with UCSF ChimeraX^59^.

### Van der Waals force analyses between PSI core and FCPIs, and within FCPIs

The van der Waals (VDW) contact analysis according to the previous method[], set the parameters in the contact function of the ChimeraX version 1.7, the VDW forces inside the FCPI subunits and all PSI core were hidden, respectively. Only the VDW forces formed by the adjacent FCPI subunits and FCPI with PSI core were shown (Fig. 5a).

### Computational analysis of Förster resonance energy transfer

The FRET rate constant is an estimated energy transfer rate constant between Chl pairs using dipole approximation to treat each Chl. The equation of FRET is $ k_{FRET} = \frac{C\kappa^2}{n^4R^6}$, where C is the spectral overlap between the electron donor’s fluorescence spectrum and the acceptor’s absorption spectrum. Empirical values of C = 32.26(CHLA to CHLA), 1.11(CHLA to CHLB), 9.61(CHLB to CHLA) and 14.45(CHLB to CHLB) yielded by Gradinaru et al was used in this calculation.

$\kappa$ is the dipole orientation factor, defined as $\kappa^2 =[\hat{\textbf{u}_D}\cdot\hat{\textbf{u}_A}-3(\hat{\textbf{u}_D}\cdot\hat{\textbf {R}_{DA}})(\hat{\textbf{u}_A}\cdot\hat{\textbf{R}_{DA}})]^2$. Where $\hat{\tex tbf{u}_D}$ and $\hat{\textbf{u}_A}$ are unit vector of displacement vector from NB to ND atom in Chl, representing the direction of dipole moment of the approximated dipole,$\hat{\textbf{R}_{DA}}$ is the unit vector of displacement vector from magnesium atom in donor Chl to magnesium atom in acceptor Chl. n stands for the refractive index, and an estimated value of n = 1.55 was taken from Gradinaru et al. R is the magnesium-to-magnesium distance between donor and acceptor Chls. The lifetime t is taken as $t_{FRET} = \frac{1}{k_{FRET}}$.

### Data availability

The cryo-EM map and atomic coordinate for the PSI-FCPI-L have been deposited in the Electron Microscopy Data Bank and the PDB (EMD-60032 and PDB ID code 8ZEH). The cryo-EM map and atomic coordinate for the PSI-FCPI-S have been deposited in the Electron Microscopy Data Bank and the PDB (EMD-60044 and PDB ID code 8ZET). The correspondence of pigment numbers in the text and renumbered in the PDB data bank (PDB: 8ZEH) was labeled in Supplementary Table 6. The data that supports the findings of this study are presented in the paper and/or the Supplementary Information.

## Acknowledgments

We thank J. Duan in the National Center for Protein Science, Shanghai Advanced Research Institute for collecting cryo-EM data. We thank Y. Lu in the College of Oceanology, Hainan University for the sequence analysis between HLR1 in N. oceanica with diatom. We thank W. Tang, L. Wang and Y. Yin of the Institute of Botany, CAS for instrumental support in sample purification, fluorescence measurement and high-performance liquid chromatography analysis, and X. Meng in the Center of Biomedical Analysis, Tsinghua University, for protein analysis.

## Author contributions

W.W. and Y.F. conceived the project. Y.F. purified the PSI-FCPI supercomplex, made biochemical characterization, and prepared figures. L.S. and X.Li collected cryo-EM images. Z.L. processed the cryo-EM data and reconstructed the final EM maps. Y.F. built the structural models. Z.L. and Y.F. refined the final models. Y.Y. helped in sequence analysis. X.Z. computes the FRET pathways. X. Liu and J. Z. helped collect cryo-EM images; F.R. and Y.W. helped cultivate diatom cells; W.W. and Y.F. wrote the manuscript; J.-R.S. modified the manuscript. Y.F., W.W., and J.-R.S. and all authors contributed to the discussion and improvement of this manuscript.

