## Supplementary material for "Structures of PSI-FCPI from *Thalassiosira pseudonana* in high light provide convergent evolution and light-adaptive strategies in diatom FCPIs": Suppl Figures and tables

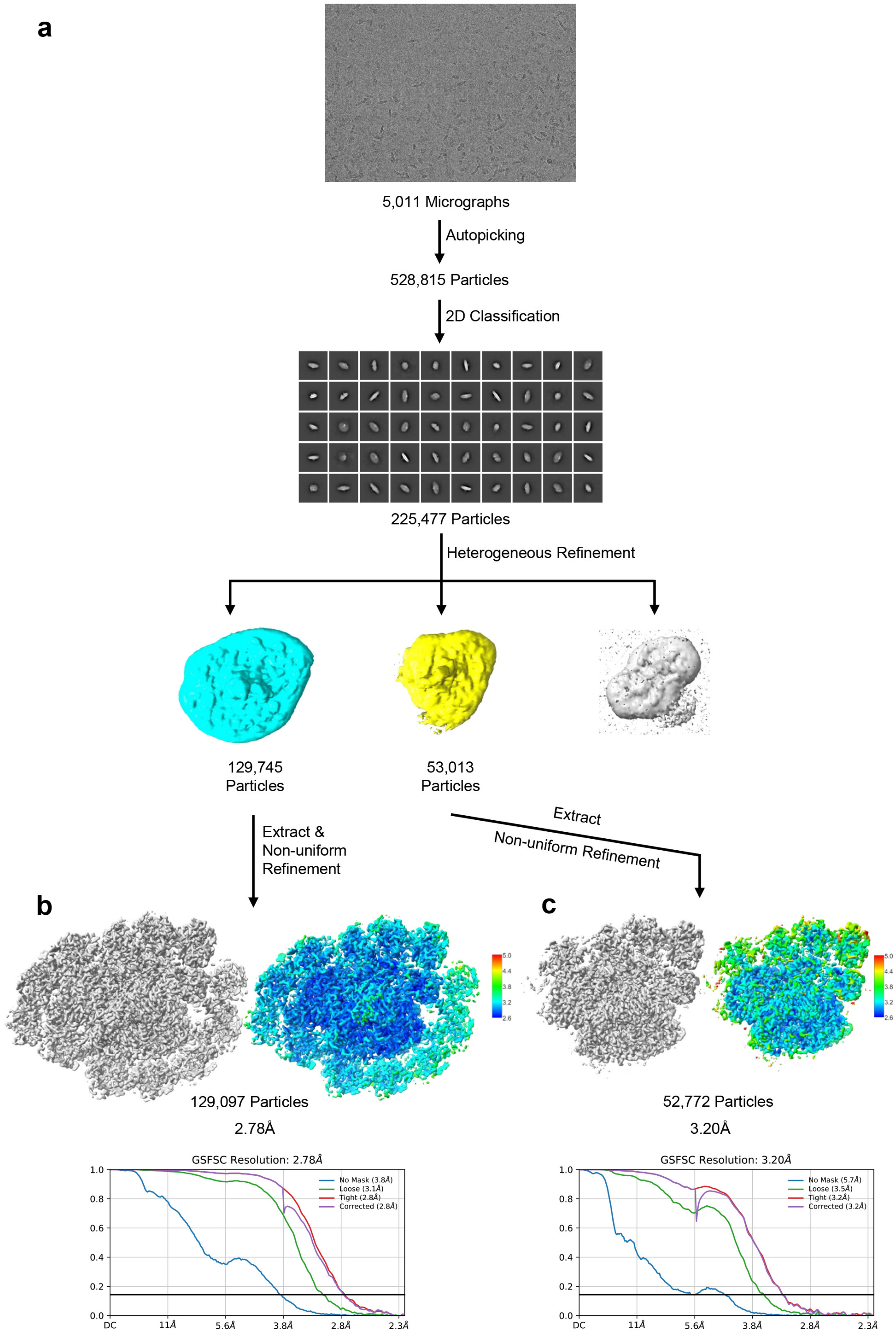


**Supplementary Fig. 1. Single particle analysis of Tp-PSI-FCPI-L and Tp-PSI-FCPI-S under high light conditions from *T. pseudonana*. a**, Flow chart for the single particle Cryo-EM analysis of Tp-PSI-FCPI-L and Tp-PSI-FCPI-S. **b-c**, Resolutions (Å) of the global and local maps and the gold standard FSC curves of the final 3D reconstruction for the global and local maps of PSI-FCPI-L and PSI-FCPI-S, respectively.

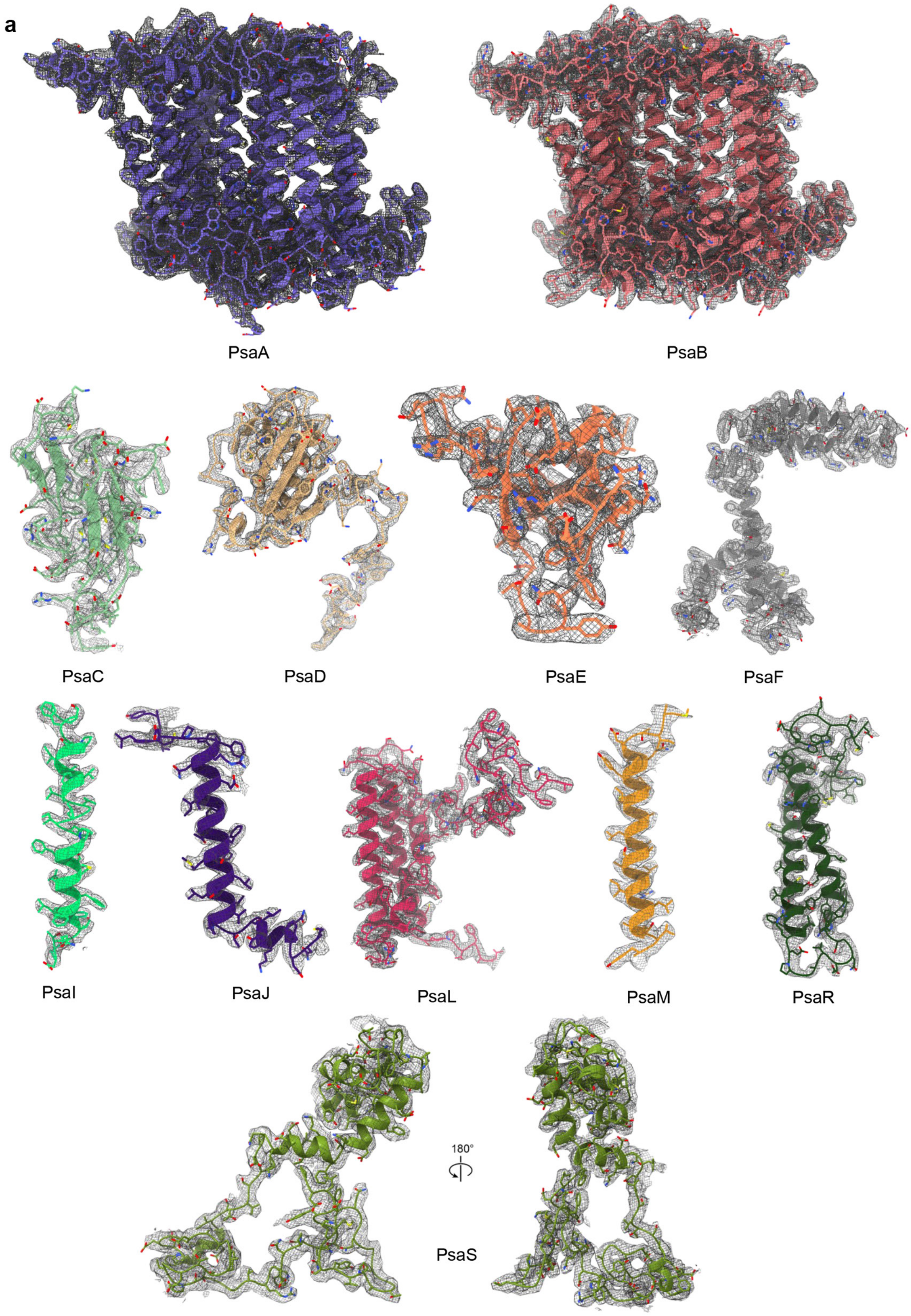
a
PsaA
PsaB
PsaC
PsaD
PsaE
PsaF
PsaI
PsaJ
PsaL
PsaM
PsaR
180°
PsaS

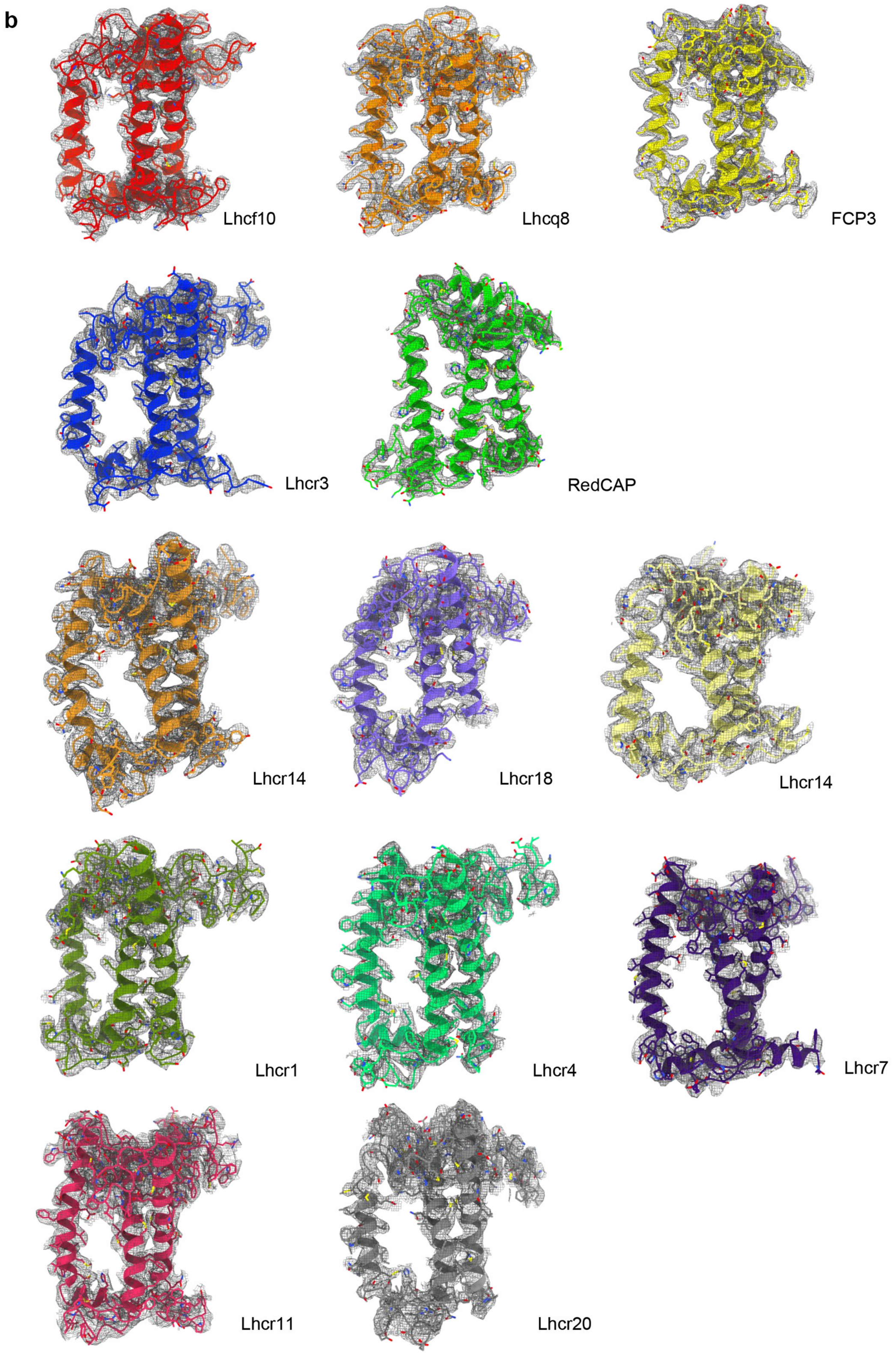
b
Lhcf10
Lhcq8
FCP3
Lhcr3
RedCAP
Lhcr14
Lhcr18
Lhcr14
Lhcr1
Lhcr4
Lhcr7
Lhcr11
Lhcr20

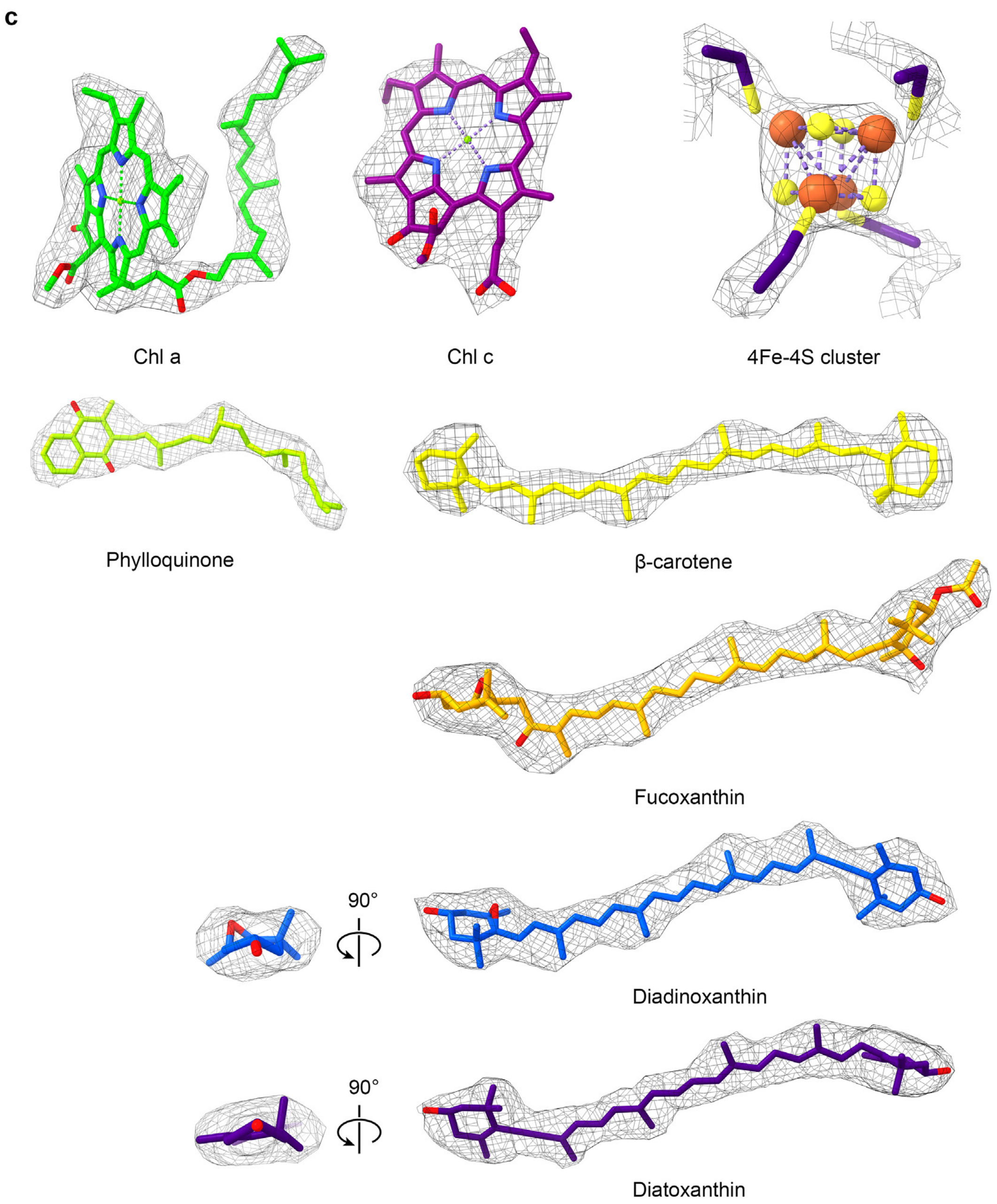


**Supplementary Fig. 2. Cryo-EM density maps for the typical PSI core subunits, antennas, pigment molecules and ligands. a-b**, The PSI core and FCPI subunits are shown in cartoons and atoms and colored as Fig. 2 and Fig. 3, the N and O atoms are colored blue and red, respectively. The density maps are shown with a threshold of 0.14 contour level (step 1) by Volume Viewer in Chimera X. **c**, Cryo-EM densities of the typical pigments and ligands in the PSI-FCPI supercomplex from *T. pseudonana*.

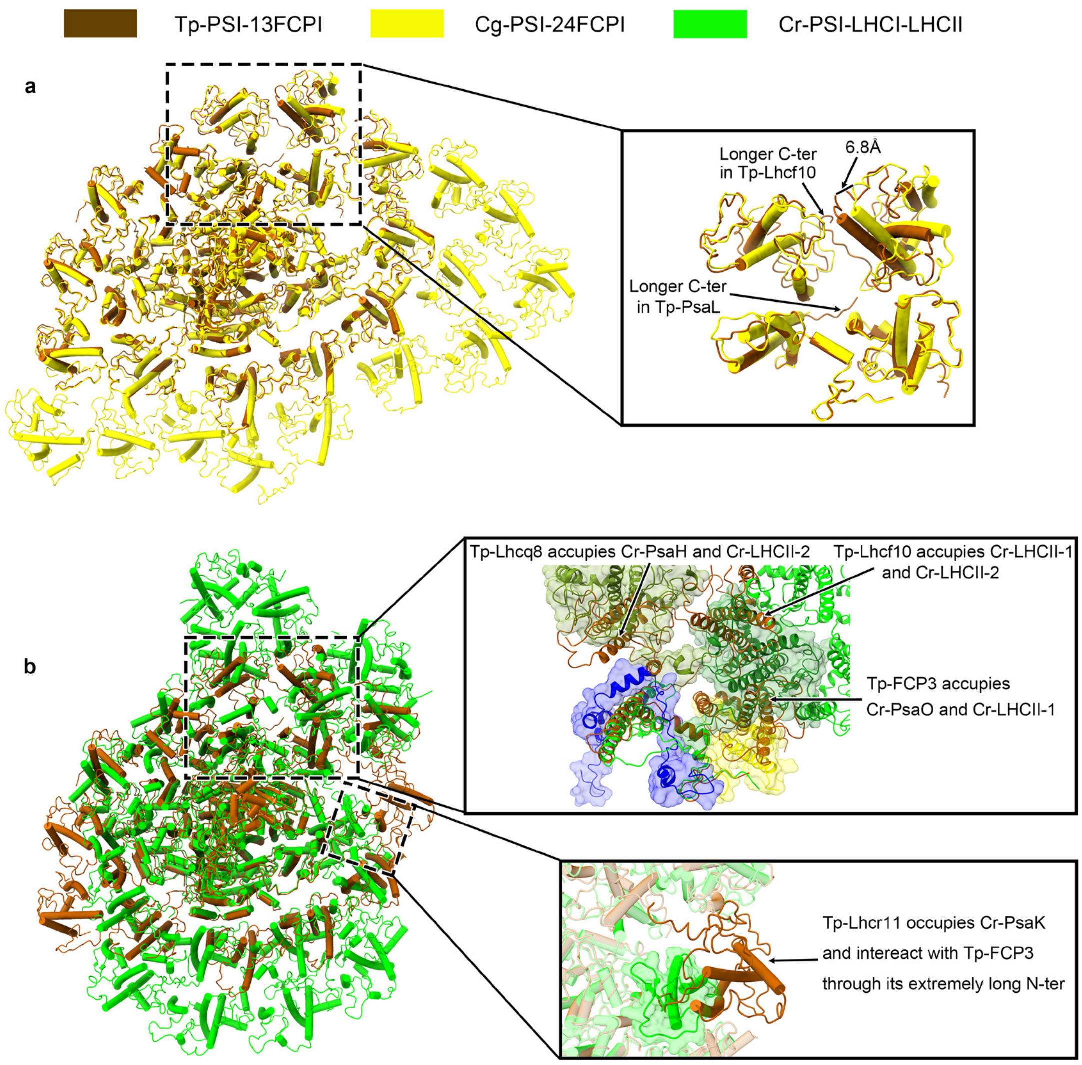


**Supplementary Fig. 3. Comparison of the Tp-PSI-FCPI-L structure with those of Cg-PSI-24FCPI and green algal PSI–LHCI–LHCII. a**, Comparison of the overall structure of Tp-PSI-PSI-FCPI-L (PDB code: 8ZEH, brown) with the huge Cg-PSI-24FCPI (PDB code 6LY5, yellow), viewed along the membrane normal to the stromal side. **b**, Comparison of the overall structure of Tp-PSI-PSI-FCPI-L (PDB code: 8ZEH, brown) with the PSI-LHCI-LHCII from *Chlamydomonas reinhardtii* (PDB code 7DZ7, yellow), viewed along the membrane normal to the stromal side.

**a**

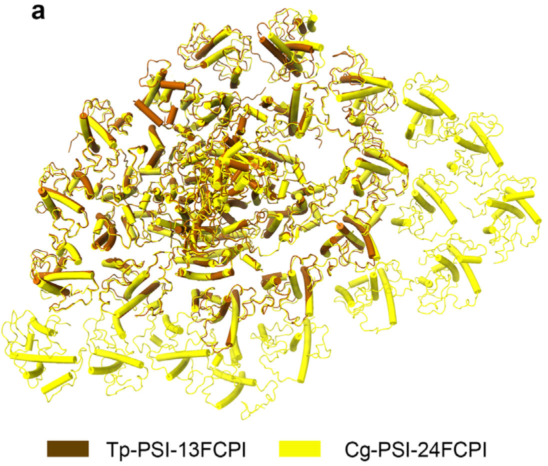


**b**

| FCPI in *T. pseudonana* | Related FCPI in *C. gracilis* |
|---|---|
| Tp-Lhcf10 | Cg-Lhcf3 |
| Tp-Lhcq8 | Cg-Lhcq12 |
| Tp-FCP3 | Cg-Lhcr9 |
| Tp-RedCAP | Cg-RedCAP |
| Tp-Lhcr3 | Cg-Lhcr1 |
| Tp-Lhcr14 | Cg-Lhcr2 |
| Tp-Lhcr18 | Cg-Lhcr3 |
| Tp-Lhcr10 | Cg-Lhcr4 |
| Tp-Lhcr1 | Cg-Lhcr5 |
| Tp-Lhcr4 | Cg-Lhcr6 |
| Tp-Lhcr7 | Cg-Lhcr7 |
| Tp-Lhcr11 | Cg-Lhcr8 |
| Tp-Lhcr20 | Cg-Lhcr10 |

**c**

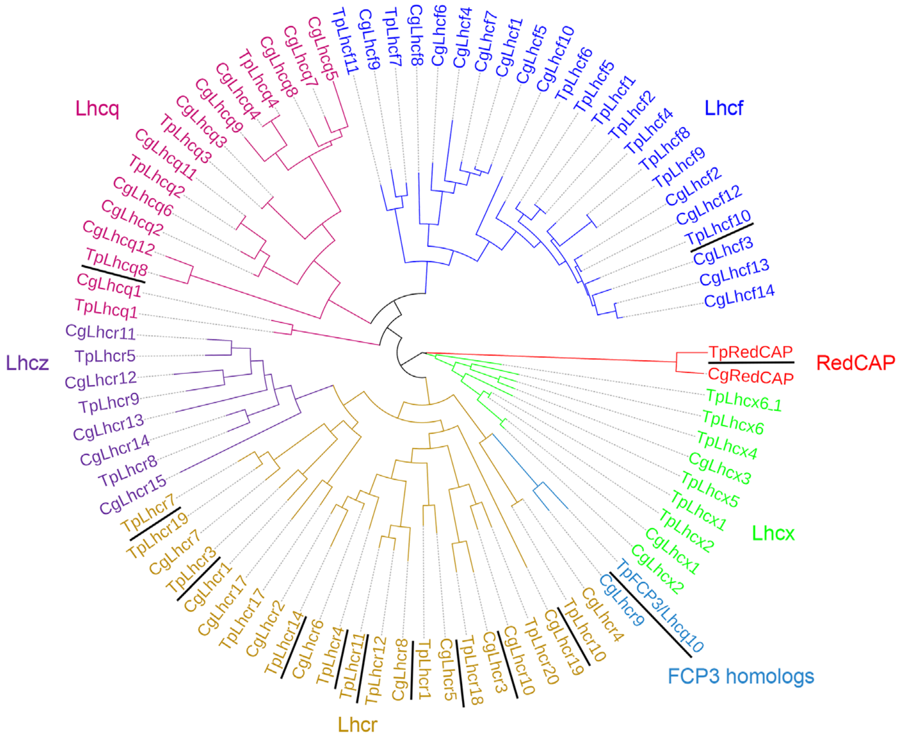


**Supplementary Fig. 4. Phylogenetic analysis and correlation of FCPIs in PSI-FCPI structures different diatoms. a**, Related structural comparisons between Tp-FCPIs (brown) and Cg-FCPIs (yellow). **b**, Correlation of the FCPI names in the homologous locations of PSI-FCPI from *T. pseudonana* and *C. gracilis*. **c**, Phylogenetic tree of typical FCPs including most Lhcf, Lhcr, Lhcx, Lhcq, Lhcz and unique RedCAP from *T. pseudonana* and *C. gracilis*, the FCPIs found in this study were underlined. Most FCP sequences of *T. pseudonana* originate from the genome information (Armbrust et al. 2004), the Cg-FCPIs originate from the previous study (Kumazawa et al. 2022), and the Cg-RedCAP originate from another study (Kato et al. 2024). The phylogenetic tree was built by Multiple Sequence Alignment of CLUSTALW, and the tree was visualized by iTOL v6. The colors of diverse clades are as follows: red, RedCAP homologs; blue, Lhcf homologs; magenta, Lhcq homologs; light purple, Lhcq homologs; dark yellow, Lhcr homologs; light blue, unique FCP3 homologs; green, unique Lhcx homologs.

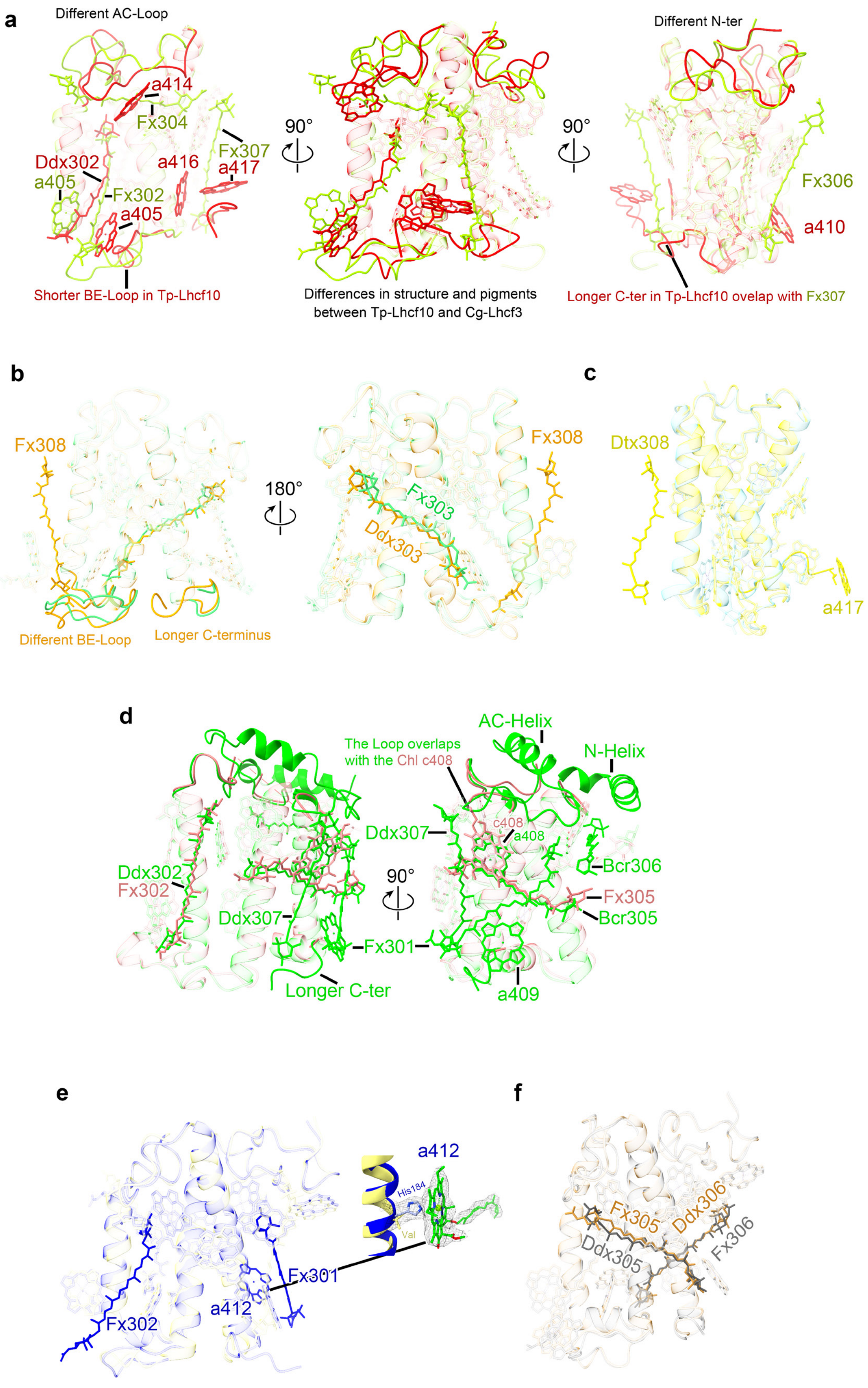
a
Different AC-Loop
a414
Fx304
Fx307
a417
Ddx302
a416
a405
Fx302
a405
Shorter BE-Loop in Tp-Lhcf10
90°
Differences in structure and pigments
between Tp-Lhcf10 and Cg-Lhcf3
90°
Different N-ter
Fx306
a410
Longer C-ter in Tp-Lhcf10 ovelap with Fx307
b
Fx308
Different BE-Loop
Longer C-terminus
180°
Fx308
Fx303
Ddx303
c
Dtx308
a417
d
The Loop overlaps
with the Chl c408
AC-Helix
N-Helix
Ddx307
c408
a408
Bcr306
Ddx302
Fx302
90°
Fx305
Ddx307
Bcr305
Fx301
Fx301
Longer C-ter
a409
e
a412
His184
Val
Fx301
a412
Fx302
f
Fx305
Ddx306
Ddx305
Fx306

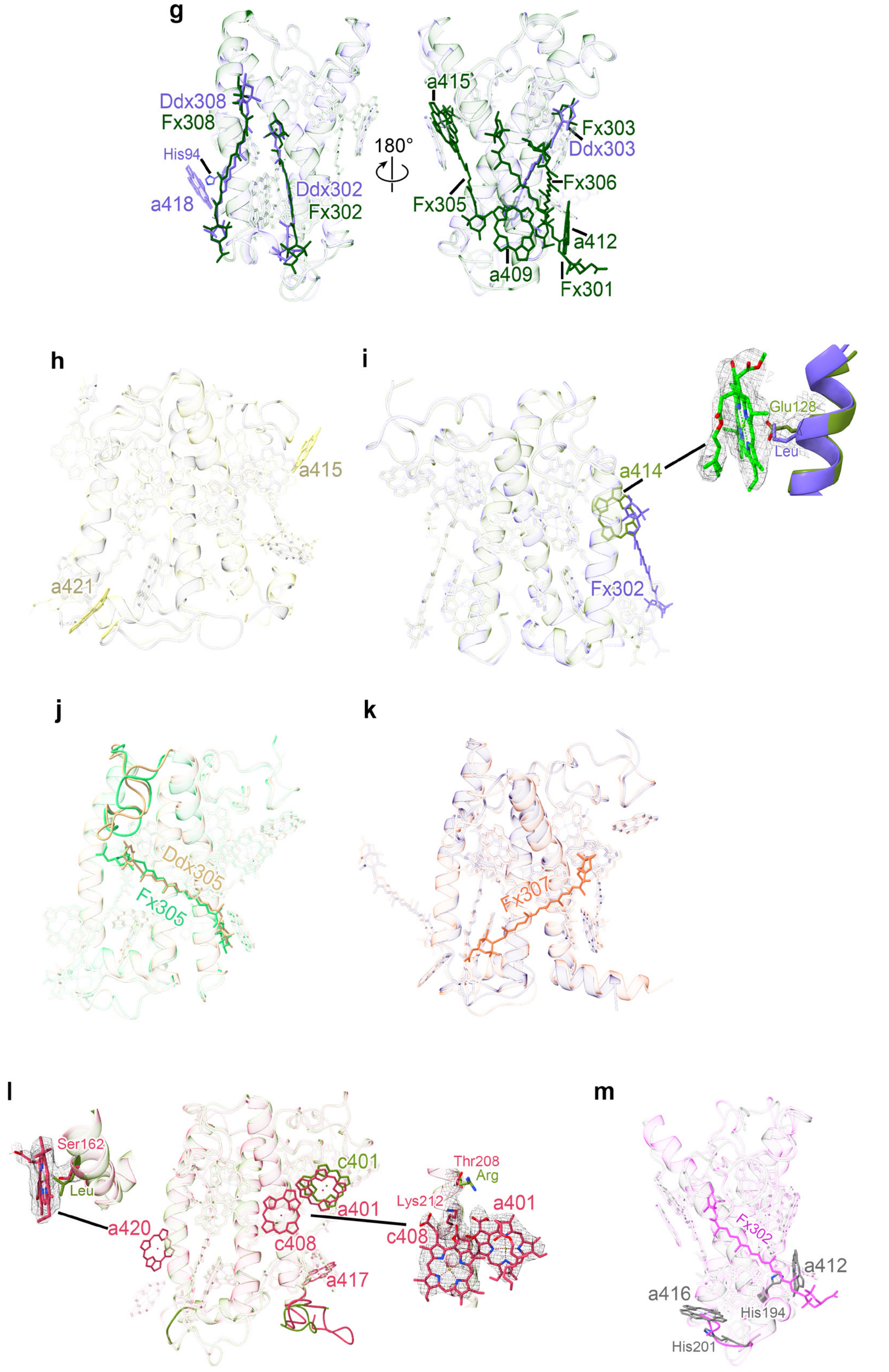


**Supplementary Fig. 5. Comparisons of structures and pigment distributions between related 13FCPIs from *T. pseudonana* and *C. gracilis*.** The conserved structures and pigment sites were transparent. The different structural areas were shown. The extra Chls or Cars of each FCPI were shown in sticks. **a**, Differences of structures and pigment distributions between structural heterogeneous Tp-Lhcf10 (red) and Cg-Lhcf3 (yellow-green) visualized in three sides. **b**, Differences of structures and pigment distributions between Tp-Lhcfq8 (orange) and Cg-Lhcq12 (light green). **c**, Differences of structures and pigment distributions between Tp-FCP3 (yellow) and Cg-Lhcr9 (cyan). **d**, Differences of structures and pigment distributions between Tp-RedCAP (green) and Cg-RedCAP (Fc13194) (light pink). **e**, Differences of structures and pigment distributions between Tp-Lhcr3 (blue) and Cg-Lhcr1 (gold). **f**, Differences of structures and pigment distributions between Tp-Lhcr14 (dark orange) and Cg-Lhcr2 (gray). **g**, Differences of structures and pigment distributions between Tp-Lhcr18 (slate blue) and Cg-Lhcr3 (forest green). **h**, Differences of structures and pigment distributions between Tp-Lhcr10 (light yellow) and Cg-Lhcr4 (light black). **i**, Differences of structures and pigment distributions between Tp-Lhcr1 (forest green) and Cg-Lhcr5 (slate blue). **j**, Differences of structures and pigment distributions between Tp-Lhcr4 (light green) and Cg-Lhcr6 (tan). **k**, Differences of structures and pigment distributions between Tp-Lhcr7 (purple) and Cg-Lhcr7 (light coral). **l**, Differences of structures and pigment distributions between Tp-Lhcr7 (deep pink) and Cg-Lhcr8 (forest green). **m**, Differences of structures and pigment distributions between Tp-Lhcr20 (gray) and Cg-Lhcr10 (magenta).

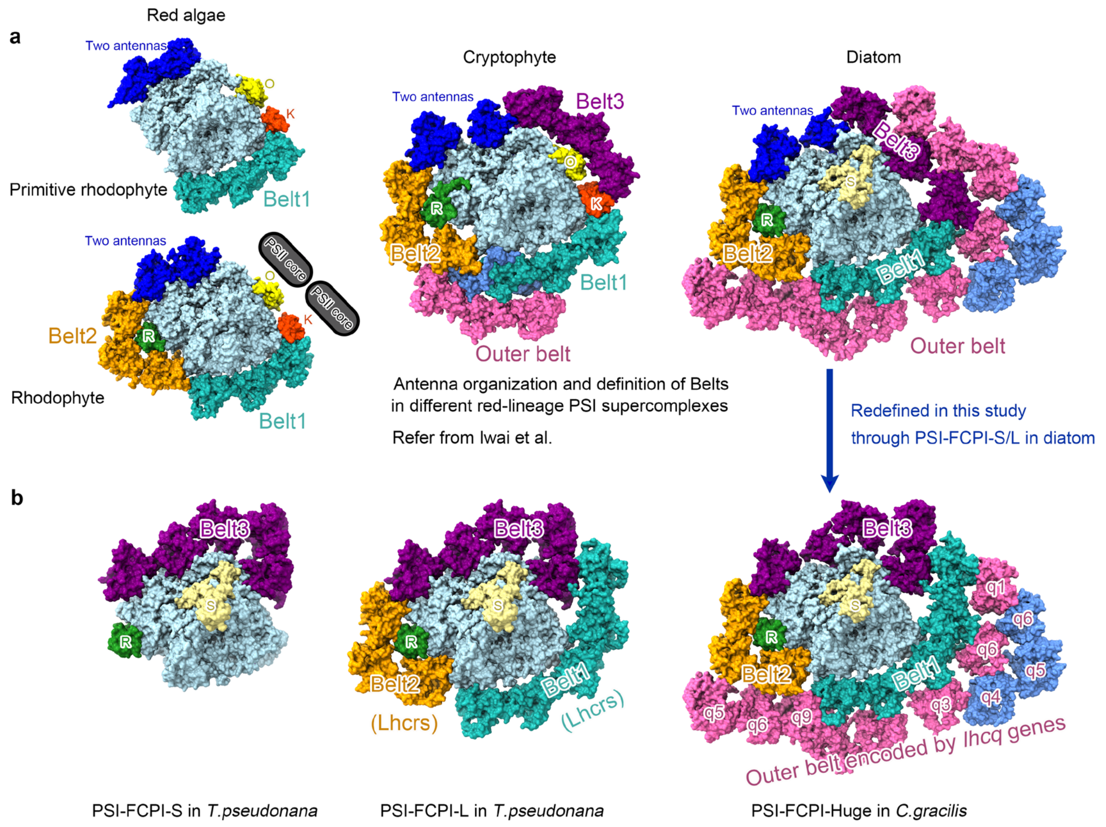


**Supplementary Fig. 6. Three FCPI belts that surround the PSI core were redefined in this study through two smaller PSI-FCPI. a**, The typical definition of the antenna belts in red lineage LHCIs, including primitive rhodophyte C. merolae (PDB: 5ZGB), rhodophyte *P. purpureum* (PDB: 7Y5E), cryptophyte *C. placoidea* (PDB: 7Y7B), and the diatom *C. gracilis* (PDB: 6LY5). **b**, The five stable FCPIs connect to the PSI core formed PSI-FCPI-S, and eight Lhcr FCPIs could flexibly connect to the PSI core formed PSI-FCPI-L. The outer belt is mainly composed of Lhcq FCPIs, modeled by PDB: 6LY5, and the FCPIs in the outer belt were named by the previous studies (Kumazawa et al. 2022) and (Calvaruso et al. 2022).

**a**

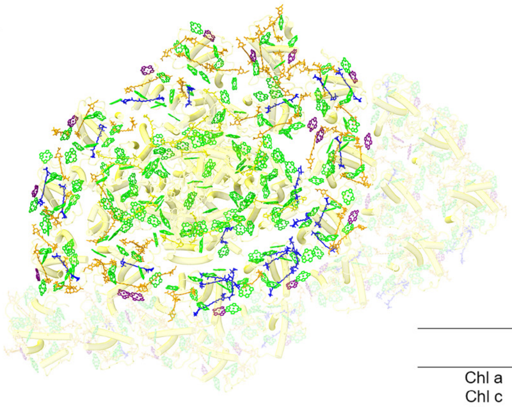

| | Total pigments in Cg-PSI-24FCPI | Pigments in homologous Cg-PSI-13FCPI | Pigments in the Outer belt |
|---|---|---|---|
| Chl a | 326 | 213 | 113 |
| Chl c | 34 | 16 | 18 |
| Bcr | 18 | 18 | 0 |
| Fx | 102 | 48 | 54 |
| Ddx/Dtx | 35 | 24 | 11 |

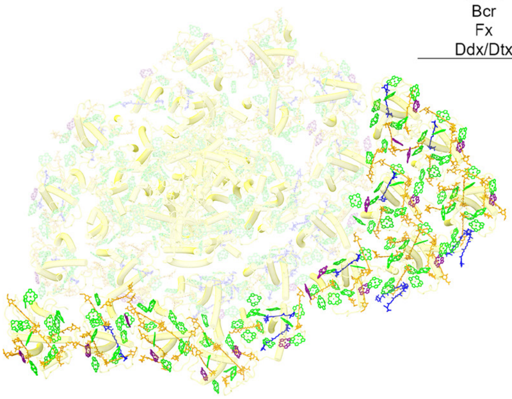

**b**

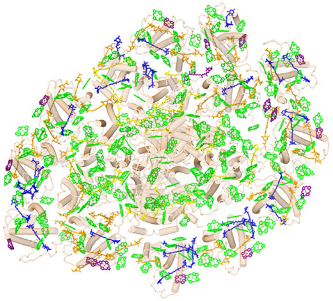

| | Total pigments in Tp-PSI-FCPI-L |
|---|---|
| Chl a | 229 |
| Chl c | 15 |
| Bcr | 21 |
| Fx | 38 |
| Ddx/Dtx | 28 |

**Supplementary Fig. 7. The number and distribution of pigments in PSI-FCPI supercomplex from *T. pseudonana* and *C. gracilis*.** **a**, The number and distribution of pigments in total Cg-PSI-24FCPI, homologous Cg-PSI-13FCPI and outer FCPI belt. **b**, The number and distribution of pigments in Tp-PSI-FCPI-L.

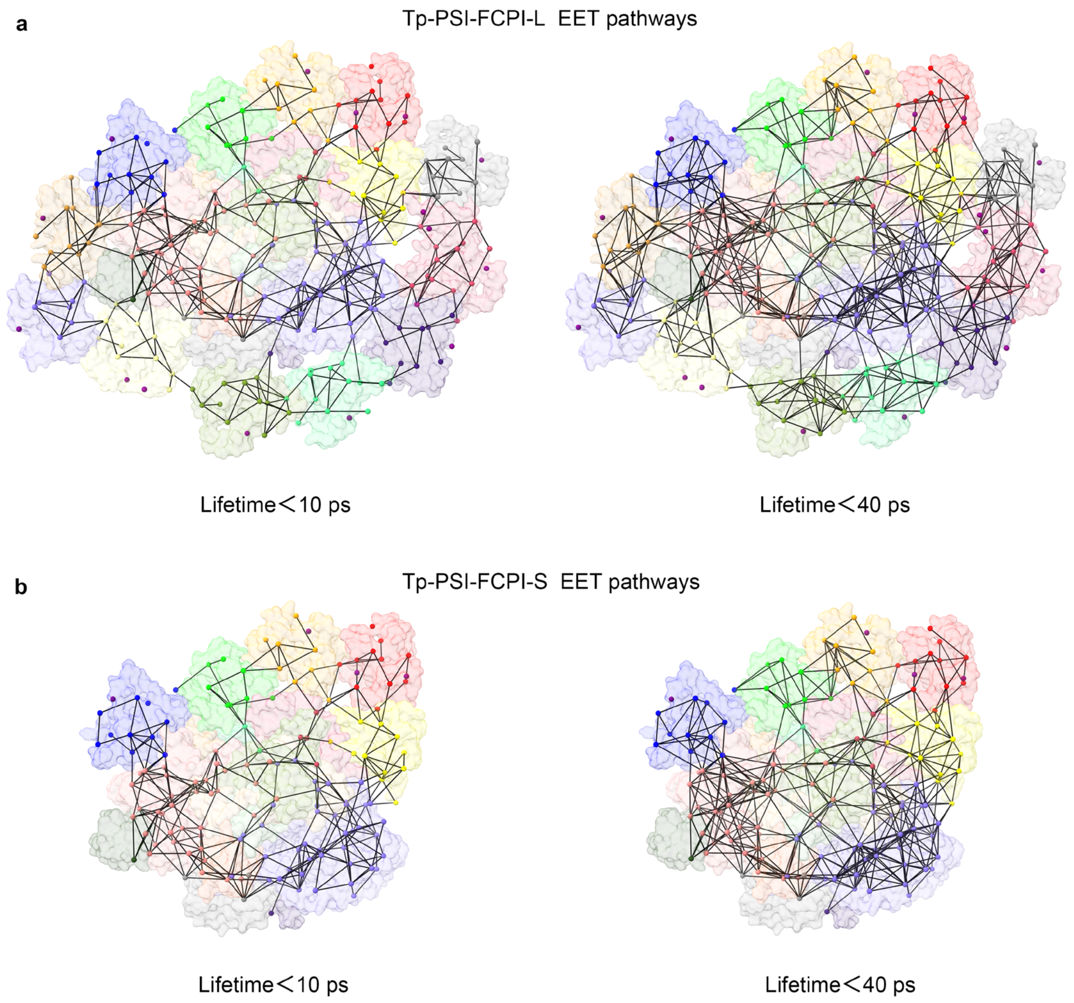


**Supplementary Fig. 8. Structure-based analysis of the FRET networks within the Tp-PSI-FCPI supercomplex.** The Chls *a* are shown in spheres and colored by different chains as in Fig.2, and all the Chls *c* are colored in purple. **a**, Organization of Chls *a* within the Tp-PSI-FCPI-L supercomplex and the FRET networks with lifetimes of less than 10 picoseconds (ps) (left) and 40 ps (right), respectively. **b**, Organization of Chls *a* within the Tp-PSI-FCPI-S supercomplex and the FRET networks with lifetimes of less than 10 ps (left) and 40 ps (right), respectively.

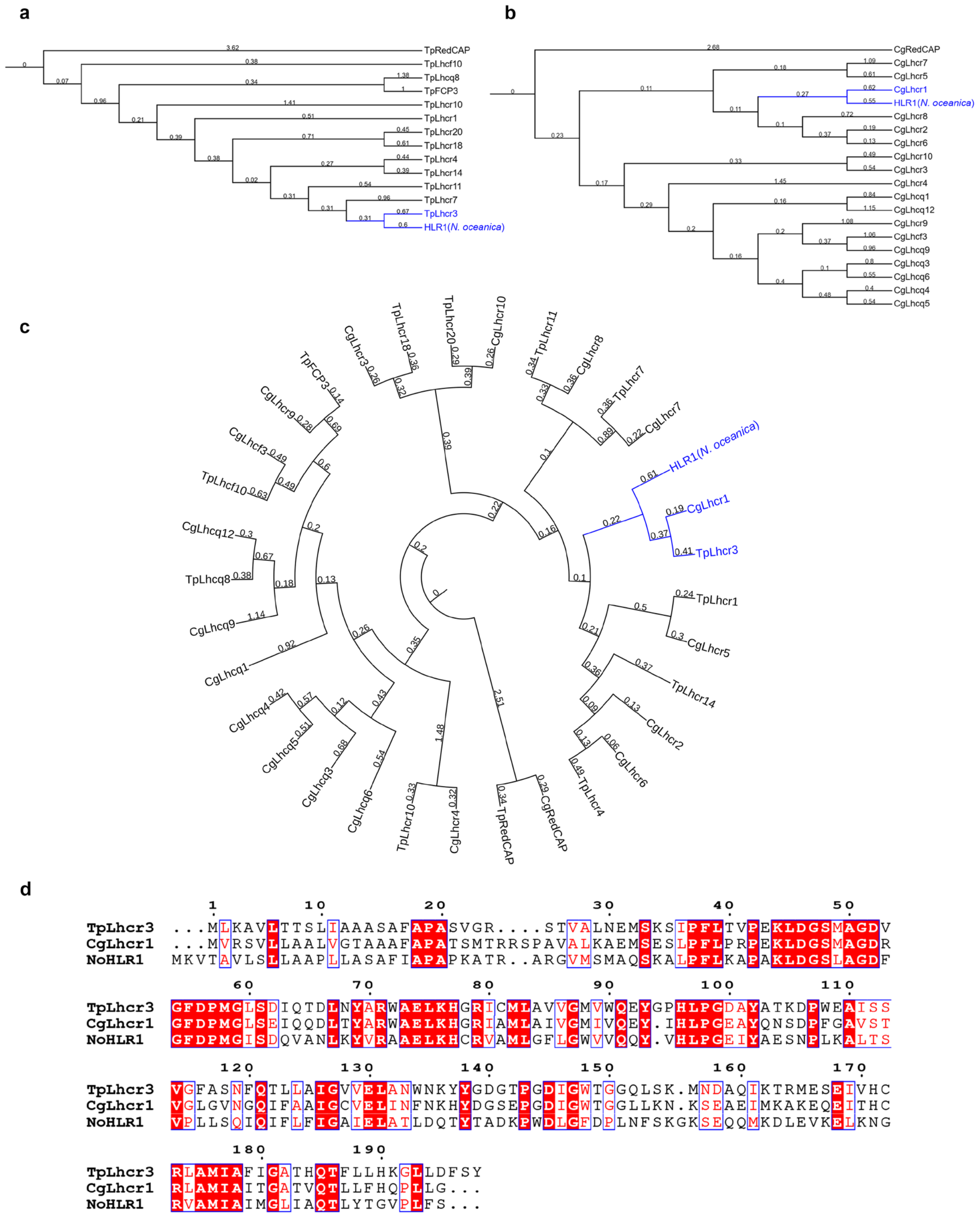


**Supplementary Fig. 9. Sequence alignment in the stramenopiles (Heterokontophyta) including homologous HLR1 from *N. oceanica* with Tp-Lhcr3 and Cg-Lhcr1 from diatoms.** **a**, Phylogenetic analysis of No-HLR1 and all the FCPIs of Tp-PSI-FCPI in this study, homologous Tp-Lhcr3 was labeled. **b**, Phylogenetic analysis of No-HLR1 and all the FCPIs of Cg-PSI-FCPI, homologous Cg-Lhcr1 was labeled. **c**, Sequence alignment of three homologous LHCR antennas. The phylogenetic tree was built by Multiple Sequence Alignment of CLUSTALW, and the tree was visualized by iTOL v6.

**Supplementary Table 1. Cryo-EM data collection, refinement, and validation statistics of PSI-FCPI-L and PSI-FCPI-S.**

| **Model statistics** | **PSI-FCPI-L** | **PSI-FCPI-S** |
|---|---|---|
| PDB code | 8ZEH | 8ZET |
| EMDB ID | EMD-60032 | EMD-60044 |
| **Data collection and processing** | | |
| Magnification | 64000× | 64000× |
| Voltage (kV) | 300 | 300 |
| Electron exposure ($e^{-}$/Å2) | 60 | 60 |
| Defocus range (μm) | -1.0 ~ -2.0 | -1.0 ~ -2.0 |
| Pixel size (Å) | 1.1 | 1.1 |
| Symmetry imposed | C1 | C1 |
| Initial particle images (no.) | 528,815 | 528,815 |
| Final particle images (no.) | 129,745 | 53,013 |
| Map resolution (Å) | 2.78 | 3.20 |
| FSC threshold | 0.143 | 0.143 |
| **Refinement** | | |
| Number of atoms | 54886 | 36581 |
| Protein residues | 4645 | 3229 |
| Ligands | 366 | 217 |
| R.m.s. deviations | | |
| Bond lengths (Å) | 0.011 | 0.012 |
| Bond angles (°) | 0.753 | 0.671 |
| Validation | | |
| MolProbity score | 2.27 | 2.19 |
| Clashscore | 12.94 | 13.61 |
| Rotamer outliers (%) | 1.14 | 0.85 |
| Ramachandran plot | | |
| Favored (%) | 93.37 | 93.72 |
| Allowed (%) | 5.68 | 5.76 |
| Disallowed (%) | 0.84 | 0.71 |

**Supplementary Table 2. Pigment-binding sites in the 13 FCPIs.** The pigment sites of the FCP dimer (Lhcf4 from *P.tricornutum*), FCP trimer (FCP05 from *C. meneghi niana*) and FCP tetramer (Lhcf1 from *C.gracilis*) were used as controls.

| | **Lhcf4 (*Pt*)** | **FCP05 (*Cm*)** | **Lhcf1 (*Cg*)** | **Lhcf10 (*Tp*)** | **Lhcq8 (*Tp*)** | **FCP3 (*Tp*)** | **RedCAP (*Tp*)** | **Lhcr3 (*Tp*)** |
|---|---|---|---|---|---|---|---|---|
| 401 | *a* | *a* | *a* | *a* | *a* | *a* | - | *a* |
| 402 | *a* | *a* | *a* | *a* | *a* | *a* | *a* | *a* |
| 403 | *c* | *c* | *c* | *c* | *a* | *a* | *a* | *a* |
| 404 | *a* | *a* | *a* | *a* | *a* | *a* | - | *a* |
| 405 | *a* | *c* | *c* | *a* | *a* | *a* | *a* | *a* |
| 406 | *a* | *a* | *c* | *a* | *a* | *a* | *a* | *a* |
| 407 | *a* | *a* | *a* | *a* | *a* | *a* | *a* | *a* |
| 408 | *c* | *c* | *c* | *c* | *c* | *a* | *a* | *c* |
| 409 | *a* | *a* | *a* | *a* | *a* | *a* | *a* | *a* |
| 410 | - | - | *a* | *a* | - | - | *a* | - |
| 411 | - | - | | - | - | - | - | - |
| 412 | - | *a* | | - | - | - | - | *a* |
| 413 | - | - | | - | - | - | - | - |
| 414 | - | - | | *a* | - | - | - | - |
| 415 | - | *a* | | - | - | - | - | *a* |
| 416 | - | - | | *a* | - | *a* | - | - |
| 417 | - | - | | *a* | - | *a* | - | - |
| 418 | - | - | | - | - | - | - | - |
| 419 | - | - | | - | - | - | - | - |
| 420 | - | - | | - | - | - | - | - |
| 421 | - | - | | - | - | - | - | - |
| 422 | - | - | | - | - | - | - | - |
| 301 | Fx | Fx | Fx | Fx | Fx | Fx | Ddx | Fx |
| 302 | Fx | Fx | Fx | Ddx | Fx | Fx | Ddx | Fx |
| 303 | Fx | Fx | Fx | Fx | Ddx | Ddx | Ddx | Ddx |
| 304 | Fx | Fx | | - | - | - | - | - |
| 305 | Fx | Fx | Fx | Fx | Fx | Fx | Ddx | Fx |
| 306 | Fx | Fx | Fx | - | - | - | Ddx | Fx |
| 307 | Fx | Fx | Fx | - | - | - | Ddx | - |
| 308 | Ddx | - | | - | Fx | Dtx | Dtx | - |
| 309 | - | - | | - | - | - | - | - |

| | **Lhcr14 (*Tp*)** | **Lhcr18 (*Tp*)** | **Lhcr10 (*Tp*)** | **Lhcr1 (*Tp*)** | **Lhcr4 (*Tp*)** | **Lhcr7 (*Tp*)** | **Lhcr11 (*Tp*)** | **Lhcr20 (*Tp*)** |
|---|---|---|---|---|---|---|---|---|
| 401 | *a* | - | *c* | *a* | *a* | *c* | *a* | *a* |
| 402 | *a* | *a* | *a* | *a* | *a* | *a* | *a* | *a* |
| 403 | *a* | *a* | *a* | *a* | *a* | *a* | *a* | *a* |
| 404 | *a* | *a* | *a* | *a* | *a* | *a* | *a* | *a* |
| 405 | *a* | *a* | *a* | *a* | *a* | *a* | *a* | *a* |
| 406 | *a* | *a* | *a* | *a* | *a* | *a* | *a* | *a* |
| 407 | *a* | *a* | *a* | *a* | *a* | *a* | *a* | *a* |
| 408 | *c* | *c* | *c* | *c* | *c* | *a* | *c* | *c* |
| 409 | *a* | - | *a* | *a* | *a* | *a* | *a* | *a* |
| 410 | *a* | *a* | - | - | *a* | - | *a* | *a* |
| 411 | - | - | - | - | - | - | - | - |
| 412 | *a* | - | - | *a* | *a* | - | - | *a* |
| 413 | - | - | - | - | - | - | - | - |
| 414 | - | - | *a* | *a* | - | - | *a* | - |
| 415 | *a* | - | *a* | *a* | *a* | *a* | *a* | - |
| 416 | - | - | - | - | - | - | *a* | *a* |
| 417 | - | - | - | - | - | - | *a* | - |
| 418 | - | *a* | - | - | - | - | - | - |
| 419 | - | - | - | - | - | - | *c* | - |
| 420 | - | - | - | - | - | - | *a* | - |
| 421 | *a* | - | *a* | - | *a* | - | - | - |
| 422 | - | - | - | - | - | - | *c* | - |
| 301 | - | - | Fx | Ddx | - | Fx | - | - |
| 302 | Fx | Ddx | Fx | Fx | Fx | Ddx | Fx | Fx |
| 303 | Ddx | Ddx | Ddx | Ddx | Ddx | Ddx | Ddx | Ddx |
| 304 | - | - | Fx | - | - | - | Fx | - |
| 305 | Fx | - | Fx | Ddx | Fx | Fx | Fx | Ddx |
| 306 | Ddx | - | Fx | Ddx | Ddx | Fx | Fx | Fx |
| 307 | - | - | Ddx | - | - | - | - | - |
| 308 | - | Ddx | - | - | - | Ddx | - | Ddx |
| 309 | - | - | Fx | - | - | - | - | - |

**Supplementary Table 3. Lengths of amino acid residues and cofactors constructed in the structures of Tp-PSI-FCPI-S and Tp-PSI-FCPI-L.**

| Protein | Source (NCBI Sequence) | Residues | Pigments | Lipids and ligands |
|---|---|---|---|---|
| PsaA | YP_874490.1 | 10-752 | 43 Chls *a*, 6 Bcr | 1 phylloquinone, 1MGDG<br>4 PG, 1 $Fe_4S_4$, 1 SQDG |
| PsaB | YP_874491.1 | 2-733 | 40 Chls *a*, 6 Bcr | 1 phylloquinone, 2 PG<br>3 DGDG |
| PsaC | YP_874576.1 | 2-81 | | 2 $Fe_4S_4$ |
| PsaD | YP_874547.1 | 8-139 | | |
| PsaE | NC_008589.1 | 1-62 | | |
| PsaF | YP_874562.1 | 25-184 | 3 Chls *a*, 1 Bcr | 1 MGDG, 1PG |
| PsaI | YP_874541.1 | 2-44 | 2 Chls *a*, 1 Bcr | |
| PsaJ | YP_874563.1 | 1-40 | 1 Chl *a*, 1 Bcr | |
| PsaL | YP_874557.1 | 2-147 | 4 Chls *a*, 3 Bcrs | 1 MGDG |
| PsaM | YP_874533.1 | 1-29 | 1 Bcr | |
| PsaR | XP_002286359.1 | 51-139 | 1 Chl *a*, 2 Fxs | |
| PsaS | XP_002287386.1 | 47-177 | | |
| Lhcf10 | XP_002295619.1 | 32-201 | 11 Chls *a*, 2 Chls *c*<br>3 Fxs, 1 Ddx | 1 MGDG |
| Lhcq8 | XP_002292077.1 | 30-194 | 8 Chls *a*, 1 Chl *c*<br>4 Fxs, 1 Ddx | 1 MGDG, 1 PG |
| FCP3 | XP_002294580.1 | 31-194 | 11 Chls *a*, 3 Fxs,<br>1 Ddx, 1 Dtx | |
| RedCAP | WWT48828.1 | 37-221 | 8 Chls *a*, 2 Bcrs<br>1 Fx, 4 Ddxs | 2 MGDG, 1PG |
| Lhcr3 | XP_002290650.1 | 31-198 | 10 Chls *a*, 1 Chl *c*<br>4 Fxs, 1 Ddx | 1 PG |
| PSI-FCPI-S | | | 142 Chls *a*, 4 Chls *c*<br>21 Bcrs, 17 Fxs<br>8 Ddxs, 1 Dtx | 2 phylloquinones, 3 $Fe_4S_4$<br>7 MGDG, 10 PG<br>1 SQDG, 1 DGDG |
| Lhcr14 | XP_002289542.1 | 33-203 | 12 Chls *a*, 1 Chl *c*<br>2 Fxs, 2 Ddxs | 2 MGDG, 1 PG |
| Lhcr18 | XP_002297322.1 | 9-161 | 8 Chls *a*, 1 Chl *c*, 3 Ddxs | 3 MGDG |
| Lhcr10 | XP_002290280.1 | 45-220 | 10 Chls *a*, 2 Chls *c*<br>7 Fxs, 1 Ddx | 1 PG |
| Lhcr1 | XP_002292353.1 | 31-204 | 11 Chls *a*, 1 Chl *c*<br>1 Fx, 4 Ddxs | |
| Lhcr4 | XP_002289997.1 | 31-202 | 12 Chls *a*, 1 Chl *c*<br>2 Fxs, 2 Ddxs | 1 MGDG, 2 PG |
| Lhcr7 | XP_002288517.1 | 32-214 | 9 Chls *a*, 1 Chl *c*<br>3 Fxs, 3 Ddxs | 1 PG |
| Lhcr11 | XP_002287251.1 | 33-249 | 14 Chls *a*, 3 Chls *c*<br>4 Fxs, 1 Ddx | 2 MGDG |
| Lhcr20 | XP_002297311.1 | 32-201 | 11 Chls *a*, 1 Chl *c*<br>2 Fxs, 3 Ddxs | |
| PSI-FCPI-L | | | 229 Chls *a*, 15 Chls *c*<br>21 Bcrs, 38 Fxs<br>27 Ddxs, 1 Dtx | 2 phylloquinones, 3 $Fe_4S_4$<br>13 MGDG, 15 PG<br>1 SQDG, 1 DGDG |

**Chl:** chlorophyll; **Fx:** fucoxanthin; **Bcr:** β-carotene; **Ddx:** diadinoxanthin; **Dtx:** diatoxanthin;

**PG**: 1,2-Dipalmitoyl-phosphatidyl-glycerole; **MGDG**: 1,2-distearoyl-monogalactosyl-diglyceride; **SQDG**: 1,2-di-O-acyl-3-O-[6-deoxy-6-sulfo-alpha-D-glucopyranosyl]-Sn-glycerol; **DGDG**: digalactosyl diacyl glycerol.

**Supplementary Table 4. The RMSD values between FCPIs in *T. pseudonana* and related FCPI in *C. gracilis*.**

| FCPI in *T. pseudonana* | Related FCPI in *C. gracilis* | RMSD (Å) / Cα atoms |
|---|---|---|
| Tp-Lhcf10 | Cg-Lhcf3 | 3.75/159 |
| Tp-Lhcq8 | Cg-Lhcq12 | 1.64/155 |
| Tp-FCP3 | Cg-Lhcr9 | 1.44/161 |
| Tp-RedCAP | Cg-RedCAP(Fc13194) | 0.91/132 |
| Tp-Lhcr3 | Cg-Lhcr1 | 1.04/163 |
| Tp-Lhcr14 | Cg-Lhcr2 | 1.34/170 |
| Tp-Lhcr18 | Cg-Lhcr3 | 1.36/150 |
| Tp-Lhcr10 | Cg-Lhcr4 | 1.23/173 |
| Tp-Lhcr1 | Cg-Lhcr5 | 0.97/168 |
| Tp-Lhcr4 | Cg-Lhcr6 | 1.02/170 |
| Tp-Lhcr7 | Cg-Lhcr7 | 1.19/182 |
| Tp-Lhcr11 | Cg-Lhcr8 | 1.76/208 |
| Tp-Lhcr20 | Cg-Lhcr10 | 1.69/163 |

**Supplementary Table 5. Chlorophyll c-binding sites in Tp-PSI-FCPI, and the sites of surrounding alkaline amino acids that interact with polar C-17 propionic acid of Chl *c*.**

| Chlorophylls c | Antenna subunit | C-17 propionic acid ligands |
|---|---|---|
| Chl *c*403 | Tp-Lhcf10 | Lys62 |
| Chl *c*408 | Tp-Lhcf10 | Lys161 |
| Chl *c*408 | Tp-Lhcq8 | Lys162 |
| Chl *c*408 | Tp-Lhcr3 | Arg164 |
| Chl *c*408 | Tp-Lhcr14 | Lys174 |
| Chl *c*408 | Tp-Lhcr18 | Lys140 |
| Chl *c*401 | Tp-Lhcr10 | Lys189 |
| Chl *c*408 | Tp-Lhcr10 | Arg182 |
| Chl *c*408 | Tp-Lhcr1 | Lys177 |
| Chl *c*408 | Tp-Lhcr4 | Lys174 |
| Chl *c*401 | Tp-Lhcr7 | Lys179 |
| Chl *c*408 | Tp-Lhcr11 | Lys212 |
| Chl *c*419 | Tp-Lhcr11 | Met38 (S) |
| Chl *c*422 | Tp-Lhcr11 | Arg37 |
| Chl *c*408 | Tp-Lhcr20 | Lys177 |

**Supplementary Table 6. Correspondence of pigments numbering in the FCPIs that are described traditionally in this text and renumbered in the PDB data bank (PDB: 8ZEH), respectively.**

| Lhcf10 | | Lhcq8 | | FCP3 | | RedCAP | | Lhcr3 | |
|---|---|---|---|---|---|---|---|---|---|
| Text No. | PDB No. | Text No. | PDB No. | Text No. | PDB No. | Text No. | PDB No. | Text No. | PDB No. |
| 301 | 302 | 301 | 301 | 301 | 203 | 301 | 301 | 302 | 201 |
| 302 | 303 | 302 | 302 | 302 | 204 | 302 | 302 | 303 | 202 |
| 303 | 304 | 303 | 303 | 303 | 205 | 303 | 303 | 305 | 203 |
| 305 | 305 | 305 | 304 | 305 | 206 | 305 | 304 | 306 | 204 |
| 401 | 306 | 308 | 305 | 401 | 207 | 306 | 305 | 401 | 205 |
| 402 | 307 | 401 | 306 | 402 | 208 | 307 | 306 | 402 | 206 |
| 403 | 308 | 402 | 307 | 403 | 209 | 308 | 307 | 403 | 207 |
| 404 | 309 | 403 | 308 | 404 | 210 | 402 | 308 | 404 | 208 |
| 405 | 310 | 404 | 309 | 405 | 211 | 403 | 309 | 405 | 209 |
| 406 | 311 | 405 | 310 | 406 | 212 | 405 | 310 | 406 | 210 |
| 407 | 312 | 406 | 311 | 407 | 213 | 406 | 311 | 407 | 211 |
| 408 | 313 | 407 | 312 | 408 | 214 | 407 | 312 | 408 | 212 |
| 409 | 314 | 408 | 313 | 409 | 215 | 408 | 313 | 409 | 213 |
| 410 | 315 | 409 | 314 | 416 | 216 | 409 | 314 | 412 | 214 |
| 414 | 316 | 510 | 315 | 417 | 217 | 410 | 315 | 415 | 215 |
| 416 | 317 | | | | | 512 | 316 | 519 | 216 |
| 417 | 318 | | | | | 513 | 317 | | |
| 512 | 301 | | | | | | | | |
| 518 | 319 | | | | | | | | |

| Lhcr1 | | Lhcr4 | | Lhcr7 | | Lhcr11 | | Lhcr20 | |
|---|---|---|---|---|---|---|---|---|---|
| Text No. | PDB No. | Text No. | PDB No. | Text No. | PDB No. | Text No. | PDB No. | Text No. | PDB No. |
| 301 | 301 | 302 | 301 | 301 | 302 | 302 | 302 | 302 | 301 |
| 303 | 302 | 303 | 302 | 302 | 303 | 303 | 303 | 303 | 302 |
| 305 | 303 | 305 | 303 | 303 | 304 | 305 | 304 | 305 | 303 |
| 306 | 304 | 306 | 304 | 305 | 305 | 306 | 305 | 306 | 304 |
| 401 | 305 | 401 | 305 | 306 | 306 | 401 | 306 | 401 | 305 |
| 402 | 306 | 402 | 306 | 308 | 307 | 402 | 307 | 402 | 306 |
| 403 | 307 | 403 | 307 | 401 | 308 | 403 | 308 | 403 | 307 |
| 404 | 308 | 404 | 308 | 402 | 309 | 404 | 309 | 404 | 308 |
| 405 | 309 | 405 | 309 | 403 | 310 | 405 | 310 | 405 | 309 |
| 406 | 310 | 406 | 310 | 404 | 311 | 406 | 311 | 406 | 310 |
| 407 | 311 | 407 | 311 | 405 | 312 | 407 | 312 | 407 | 311 |
| 408 | 312 | 408 | 312 | 406 | 313 | 408 | 313 | 408 | 312 |
| 409 | 313 | 409 | 313 | 407 | 314 | 409 | 314 | 409 | 313 |
| 412 | 314 | 410 | 314 | 408 | 315 | 410 | 315 | 410 | 314 |
| 414 | 315 | 412 | 315 | 409 | 316 | 414 | 316 | 412 | 315 |
| 415 | 316 | 415 | 316 | 415 | 317 | 415 | 317 | 416 | 316 |
| 416 | 317 | 421 | 317 | 517 | 318 | 416 | 318 | | |
| | | 519 | 318 | 518 | 301 | 419 | 319 | | |
| | | 521 | 319 | | | 420 | 320 | | |
| | | | | | | 422 | 321 | | |
| | | | | | | 502 | 322 | | |

| Lhcr14 | | Lhcr18 | | Lhcr10 | |
|---|---|---|---|---|---|
| Text No. | PDB No. | Text No. | PDB No. | Text No. | PDB No. |
| 302 | 302 | 302 | 201 | 301 | 301 |
| 303 | 303 | 303 | 202 | 302 | 302 |
| 305 | 304 | 308 | 203 | 303 | 303 |
| 306 | 305 | 402 | 204 | 304 | 304 |
| 401 | 306 | 403 | 205 | 305 | 305 |
| 402 | 307 | 404 | 206 | 306 | 306 |
| 403 | 308 | 405 | 207 | 307 | 307 |
| 404 | 309 | 406 | 208 | 309 | 308 |
| 405 | 310 | 407 | 209 | 401 | 309 |
| 406 | 311 | 408 | 210 | 402 | 310 |
| 407 | 312 | 410 | 211 | 403 | 311 |
| 408 | 313 | 511 | 213 | 404 | 312 |
| 409 | 314 | | | 405 | 313 |
| 410 | 315 | | | 406 | 314 |
| 412 | 316 | | | 407 | 315 |
| 415 | 317 | | | 408 | 316 |
| 421 | 318 | | | 409 | 317 |
| 518 | 319 | | | 414 | 318 |
| 519 | 320 | | | 415 | 319 |
| | | | | 421 | 320 |
| | | | | 501 | 321 |
